# Population Dynamics of Immunological Synapse Formation Induced by Bispecific T-cell Engagers Predict Clinical Pharmacodynamics and Treatment Resistance

**DOI:** 10.1101/2022.04.18.488626

**Authors:** Can Liu, Jiawei Zhou, Stephan Kudlacek, Timothy Qi, Tyler Dunlap, Yanguang Cao

## Abstract

Effector T cells form immunological synapses (IS) with recognized target cells to elicit cytolytic effects. Facilitating IS formation is the principal pharmacological action of most T cell-based cancer immunotherapies. However, the dynamics of IS formation at the cell population level, the primary driver of the pharmacodynamics of many cancer immunotherapies, remains poorly defined. With classic immunotherapy CD3/CD19 bispecific T cell engager (BiTE) as a model system, we integrate experimental and theoretical approaches to investigate the population dynamics of IS formation and their relevance to clinical pharmacodynamics and treatment resistance. Our models produce experimentally consistent predictions when defining IS formation as a series of spatiotemporally coordinated events driven by molecular and cellular interactions. The models predict tumor-killing pharmacodynamics in patients and reveal trajectories of tumor evolution across anatomical sites under BiTE immunotherapy. Our model highlight the bone marrow may serve as a sanctuary site permitting tumor evolution and antigen escape. The models also suggest the optimal dosing regimens as a function of tumor growth and patient T cell abundance that confer adequate tumor control with minimal disease evolution. This work has implications for developing more effective T cell-based cancer immunotherapies.

## Introduction

Immunotherapy is a type of cancer treatment that helps patients’ immune system fight cancer, and these therapies include immune checkpoint inhibitors and bispecific T cell engagers (BiTE). Despite many successes, multiple challenges persist. For instance, only a fraction of patients respond, and many responders eventually relapse (***Nagorsen et al., 2012; Thakur et al., 2018; Topp et al., 2014***). The primary pharmacological action of cancer immunotherapies is to activate or reinvigorate the effector T cells to find and engage tumor cells and eventually form cytolytic immunological synapses (IS). BiTE, as a unique type of cancer immunotherapy, can redirect T cells to bind specific antigens on tumor cells and form IS. Facilitating the formation of IS through tight intercellular apposition lead to cytolysis of the tumor cell. IS formation is therefore a critical step for BiTE pharmacodynamics (***Nagorsen et al., 2012***). Like the IS formed between antigen-presenting cells and T cells, BiTE-induced IS formation between tumor cells and T cells is a precisely orchestrated cascade of molecular and cellular interactions (***Delon and Germain, 2000; Roda-Navarro and Álvarez-Vallina, 2020***). Understanding the dynamics of IS formation at the cell population level and factors governing this process hold promise to predicting pharmacodynamics and treatment resistance, one of the most critical issues for immunotherapies.

The molecular mechanisms of IS formation have been widely studied to identify potential molecular targets for cancer immunotherapy (***Finetti et al., 2018; Xiong et al., 2018***). However, IS formation at the macroscopic cell population level extends beyond molecular crosslinking and involves multiple steps of intercellular interactions, such as T cells scanning of target cells in the tumor microenvironment, slowing their motility upon recognition, and establishing intercellular adhesion in response to signals generated by the first encounter (***Dustin and Cooper, 2000; Fousek et al., 2021***). Previous pharmacodynamic models of BiTE immunotherapy have focused, almost exclusively, on the mechanisms of molecular crosslinking, with little consideration for macroscopic intercellular interactions (***Betts et al., 2019; Jiang et al., 2018; Schropp et al., 2019; Song et al., 2021***). We sought to investigate how intercellular interactions at the population level could provide further insight into IS formation dynamics, BiTE pharmacodynamics, and mechanisms of resistance.

Tumor cells reside in dynamic and heterogeneous microenvironments. Suboptimal BiTE efficacy could result from insufficient numbers of effector cells, poor drug penetration into tumor tissue, and antigen loss leading to immune escape. Characterizing IS formation dynamics under diverse conditions, such as varying T cell density, BiTE concentration, antigen binding affinity, and antigen expression, could provide insights into factors determining BiTE pharmacodynamics and possible mechanisms of tumor resistance. Bell. et al. developed theoretical models of cell-cell adhesion (***Bell, 1978***) that highlighted not only the importance of cell-surface receptors and ligands, but also biophysical factors including receptor diffusivity on the cell membrane and hydrodynamic shearing forces. By now, to our knowledge, there has been no theoretical framework developed to characterize the population dynamics of IS formation, limiting our ability to predict the pharmacodynamics of and resistance to cancer immunotherapies.

In this study, we applied imaging flow cytometry to quantify BiTE-induced IS formation dynamics under various experimental conditions. Additionally, we developed theoretical models to simulate IS formation on different spatiotemporal scales to interrogate factors influential to this process. After considering patient-specific parameters, models incorporating IS formation dynamics adequately predicted tumor-killing pharmacodynamics in patients. These models revealed trajectories of antigen escape and tumor evolution across anatomical sites, and predicted optimal doses and regimens that could confer effective tumor control with reduced disease evolution. Our work shows substantial implications for developing effective T cell-based cancer immunotherapies.

## Results

### Experimental design and theoretical models

The formation dynamics of the BiTE-induced IS were the focus of this study. As depicted in ***Figure 1a***, CD3^+^ Jurkat were used as effector cells (E) while CD19^+^ Raji were used as target cells (T). Jurkat and Raji cells were sorted into three subpopulations, high (H), medium (M), and low (L), based on their membrane expressions of CD3 or CD19, respectively (***Supplementary Figure S1***). Effector and target cells were then co-incubated in the presence of blinatumomab. IS formation dynamics were visualized and quantified by imaging flow cytometry (***Figure 1a, Supplementary Figure S2***). We quantified the dynamics of IS formation under various experimental conditions, including different cell densities, antigen expressions, incubation durations, E:T ratios, and antibody concentrations. Multiple types of IS were observed and quantified, including “typical” IS with one effector and one target cell (ET) and “variants” with three or more cells engaged, such as ETE, ETT, ETET.

**Figure 1.**
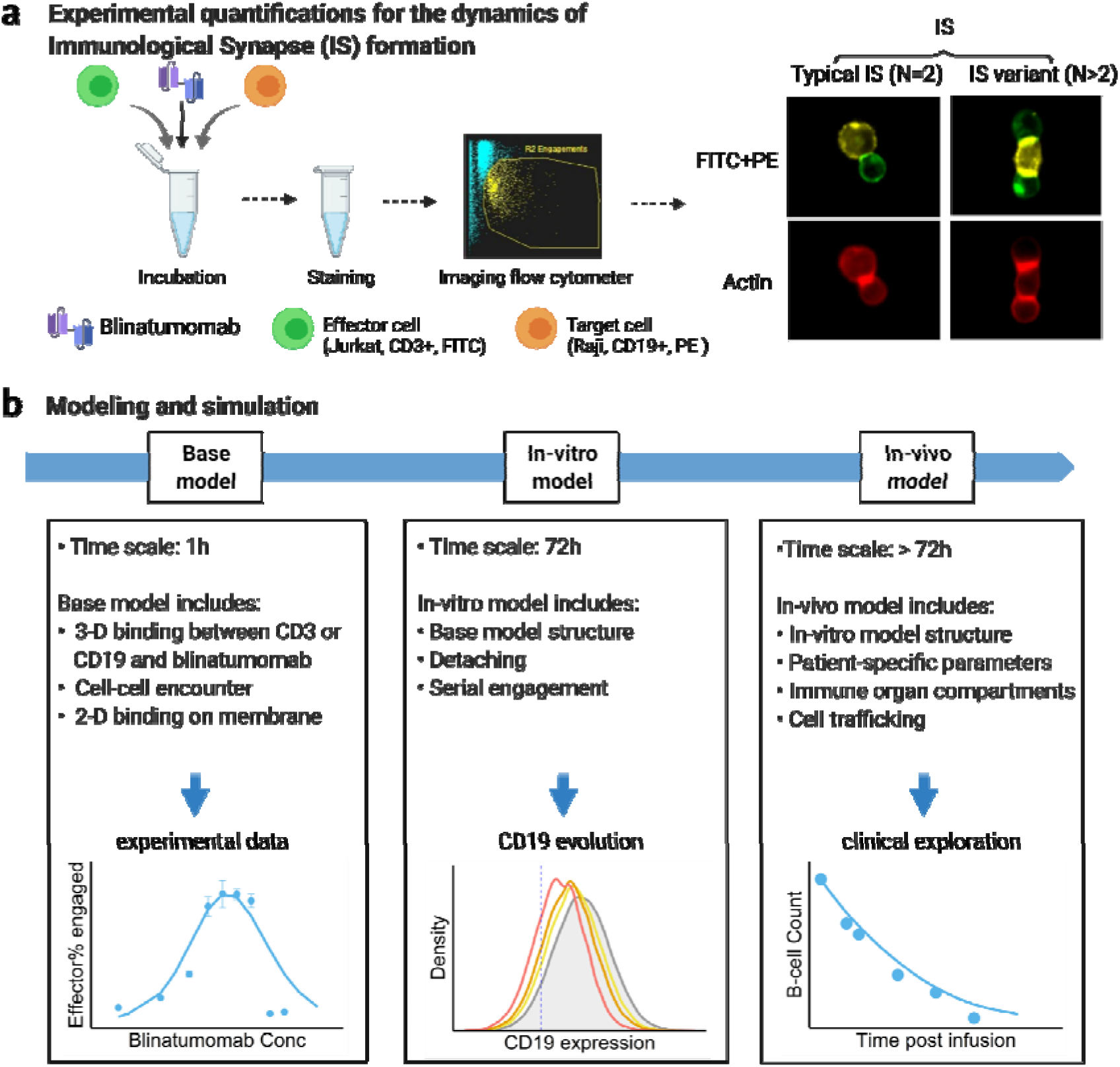
Schematic of experimental design and theoretical models. The study examined the formation dynamics of immunological synapses (IS) elicited by a bispecific T cell engager (BiTE). **a**, The abundance and dynamics of IS formation were quantified by imaging flow cytometry under various experimental conditions. **b**, Three mechanistic agent-based models were developed for comprehensive characterization of cell-cell engagement and tumor-killing effect on different spatiotemporal scales. 3-D, three-dimensional; 2-D, two-dimensional.

We developed three mechanistic agent-based models to investigate IS formation and tumor-killing on different spatiotemporal scales. These models were developed in a stepwise fashion and calibrated with experimental or clinical data. As shown in ***Figure 1b***, the base model was developed for predicting IS formation dynamics within 1 hour. The base model consists of three fundamental components during IS formation: three-dimensional (3D) binding between antibody and antigens (CD3 or CD19) to form binary complexes, the probability of cell-cell encounter, and cell-cell adhesion driven by two-dimensional (2D) binding to form ternary complexes (CD3-BiTE-CD19) on the cellular membrane. Model structure for the base model is provided in the next section.

Next, the base model was extended with serial cell-cell engagement dynamics to capture IS formation and tumor-killing up to 72 hours; we refer to this as the in-vitro model. The in-vitro model could evaluate the chance for target antigen loss (i.e., CD19), a common mechanism of immune escape and therapeutic resistance to BiTEs (***Thakur et al., 2018; Topp et al., 2014***).

Last, the in-vitro model was expanded to include cell-cell engagement in anatomically distinct compartments of the body. This model considered infiltration gradients of T and B cells and organ-to-organ cell trafficking. We refer to this expanded model as the in-vivo model. This model integrated patient-specific parameters to predict clinically observed profiles of tumor killing and relapse. The in-vivo model was also applied to support simulation of antigen escape and tumor evolution across anatomical sites and compare dosing regimens for effective tumor control in light of tumor evolution.

### Model structure (Base model)

The major mechanism of BiTE pharmacology is to produce ternary complexes (CD3-BiTE-CD19) on opposing cell surfaces, driving IS formation. The dynamics of IS formation are, essentially, two independent and indispensable cellular processes mediated by cell-cell encounter and cell-cell adhesion. Compared to molecular scale-focused pharmacodynamics models of BiTEs (***Betts et al., 2019; Jiang et al., 2018; Schropp et al., 2019; Song et al., 2021***), our models highlight the importance of these two cellular processes for IS formation in the context of macroscopic and biophysical forces (***Figure 2***).

**Figure 2.**
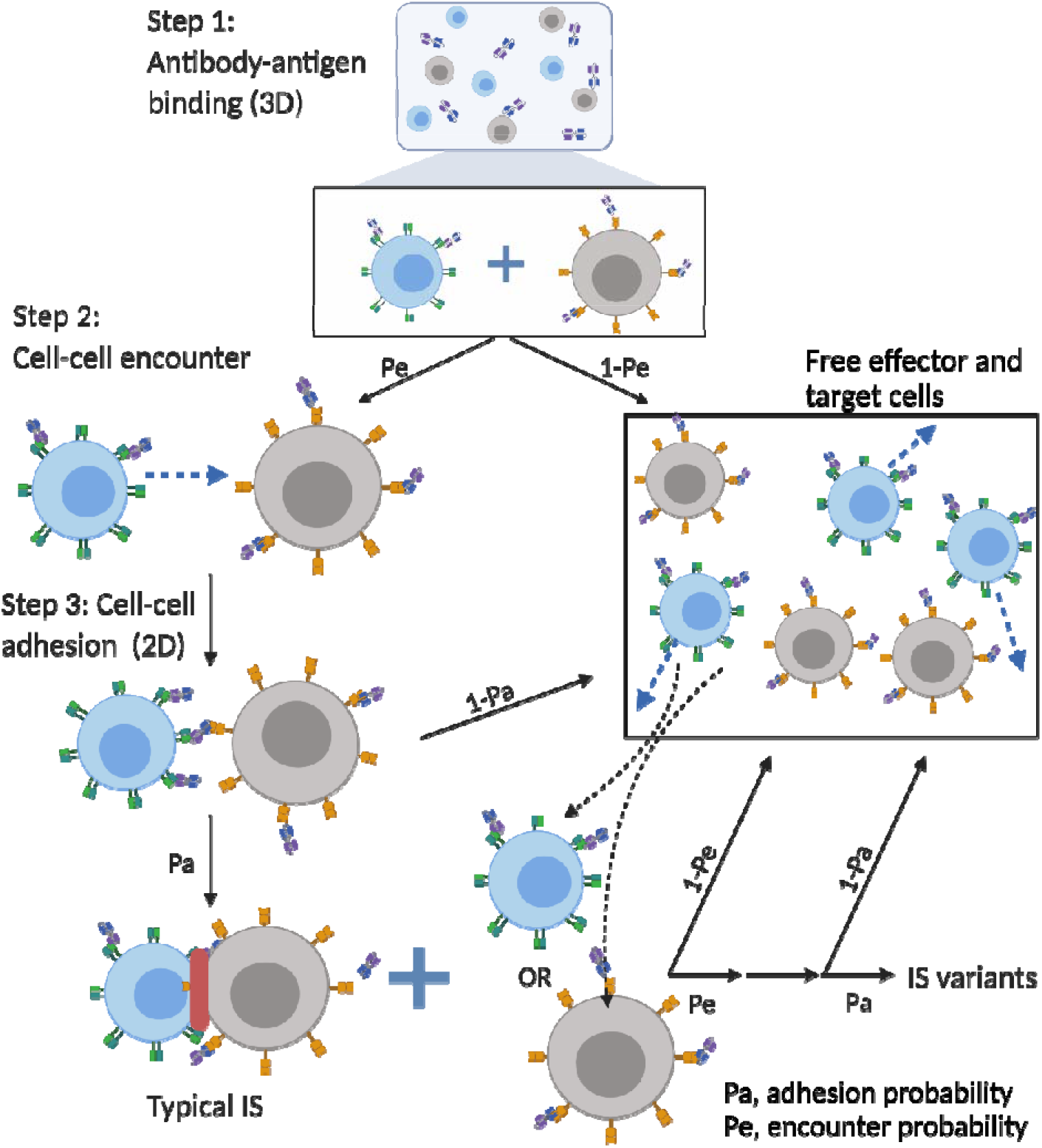
The base model included three essential steps to describe the process of IS formation induced by BiTEs. Step 1: three-dimensional (3D) antibody-antigen binding in the media to form a binary complex; Step 2: cell-cell encounter, with encounter probability (Pe) dictated by cell motility and density; Step 3: cell-cell adhesion and IS formation, with adhesion probability (Pa) driven by the density of ternary complexes formed on the cell-cell contact area (two-dimensional (2D) binding) during contact. Newly formed typical IS had a chance to engage additional free effector or target cells to form an IS variant.

In the base model (***Figure 2***), three essential steps are defined: antibody-antigen binding to form binary complexes (BiTE-CD3 and BiTE-CD19) (step 1), effector-target cell encounter probability defined as a function of cell mobility and density (step 2), and effector-target cell adhesion probability defined as a function of ternary complexes formed during contact (step 3). Model equations are provided in ***Supplementary Figure S3-S9***. Antibody-antigen binding to form binary complexes (BiTE-CD3 and BiTE-CD19) were assumed to be 3D processes that reached rapid equilibrium prior to cellular-scale events. Free effector cells were assumed to follow Brownian motion (***Celli et al., 2012***) and encounter target cells at a probability proportional to cell density and motility in the incubation environment (***Figure 2***).

After cell-cell encounter, the probability of adhesion was modeled to increase as a function of the number of ternary complexes (CD3-BiTE-CD19) formed between cells during the duration of contact. Cell-cell complexes with a high number of ternary complexes would therefore have a higher probability to adhere and eventually form IS. Ternary complex formation on opposing cells was assumed to be restricted to the cell membrane (2D binding; ***Supplementary Figure S6***). The derivation of 2D binding affinities is provided in ***Supplementary Figure S8***. Cells that failed to adhere upon encounter (futile contact) were assumed to diffuse away and become free to repeat the same process. Two cells engaged in a typical IS were also considered capable of engaging additional effector or target cells to become an IS variant, per our experimental observations (***Figure 2***). The model considered IS variants comprising up to 4 cells.

Model details and parameters for base, in-vitro and in-vivo models are provided in *Supplementary Methods*, *Supplementary Figure S3-S9*, and *Supplementary Table S1-S3*.

### Effects of BiTE concentration, cell density, and antigen expression on IS formation

IS formation dynamics were quantified by imaging flow cytometry. Representative images of non-engaged (futile contact), typical IS, and multiple IS variants are shown in ***Figure 3a***. Contact between effector and target cells was evaluated in brightfield and FITC + PE channels. Their interfaces were classified as bona fide IS when there was a high intensity of actin (red) at the contact site, as F-actin is known to polymerize and locally concentrate at sites of interface (***Dustin and Cooper, 2000***).

**Figure 3.**
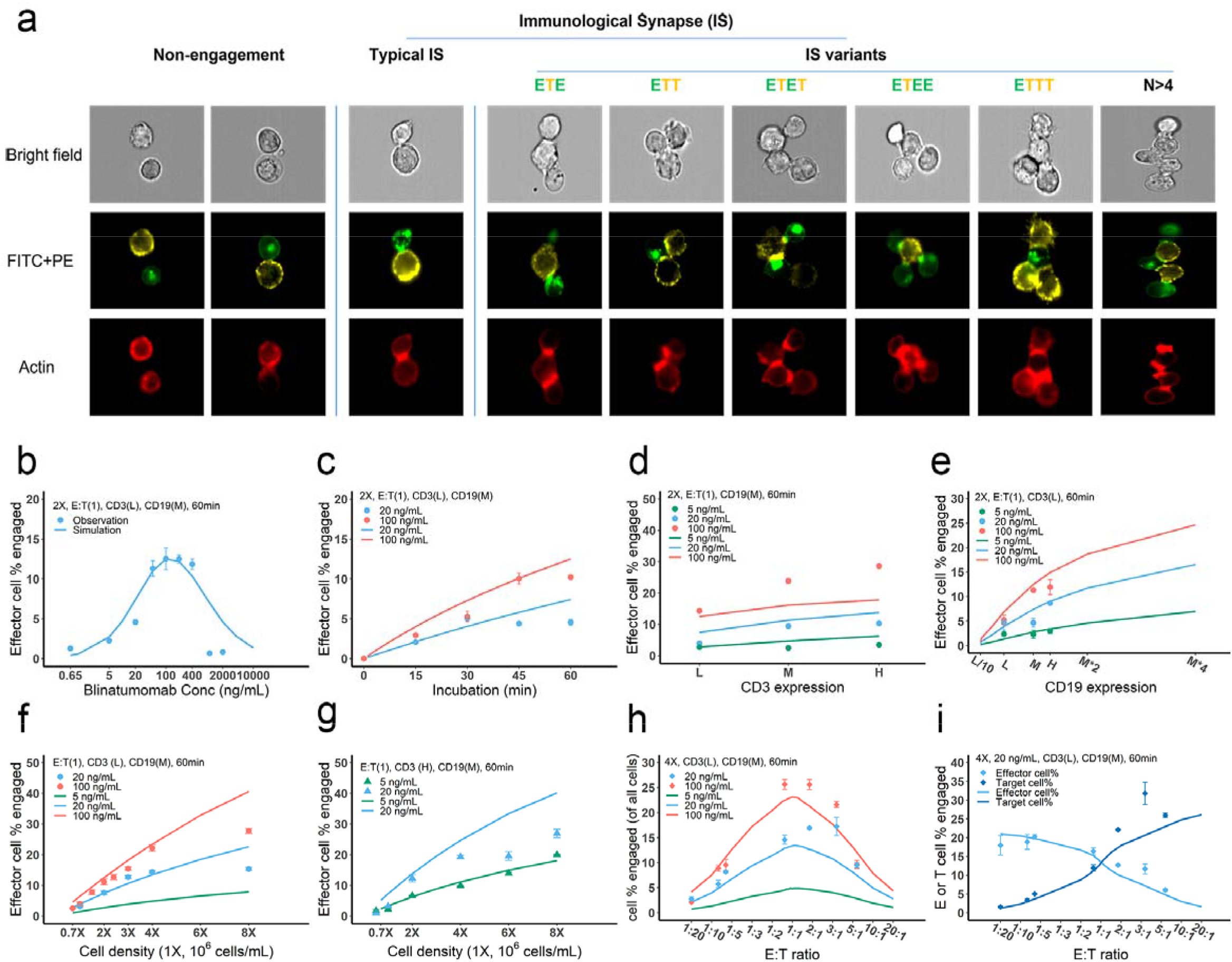
Dynamics of immunological synapse (IS) formation induced by BiTE under different conditions. **a**, Representative image of non-engagement (futile encounter), typical IS, and other IS variants. Green (FITC), effector cells (E); Yellow (PE), target cells (T). **b-i**, The effects of drug concentration (**b**), incubation duration (**c**), antigen density (**d, e**), cell density (**f, g**) and E:T ratio (**h, i**) on IS formation. The base model was applied to simulate IS formation under different conditions. Observations are dots (with SE) and model simulations are solid curves. 2X, 2×10^6^ cells/mL; E:T(1), E:T ratio = 1; CD3(L), CD3 expression (Low); CD3(H), CD3 expression (High); CD19(M), CD19 expression (medium); 5, 20, 100 ng/mL, blinatumomab concentration; 60 min, incubation duration.

To investigate the key influential factors of IS formation, we explored multiple experimental conditions by varying BiTE concentration (0.65-2000 ng/ml), incubation duration (0-60 min), antigen expression (three levels for either CD3 or CD19), cell density (0.7-8 million), and E:T ratio (0.05-6) (***Figure 3b-i***). The fraction (%) of effector cells engaged in IS was quantified to inform IS formation dynamics. We also ran these experiments virtually using the base model to test the model’s predictive performance and to explore mechanistic hypotheses.

In ***Figure 3b-i***, the observations (symbols) and model simulations (lines) overlapped, indicating good base model performance. IS formation in vitro exhibited a bell-shaped relationship to BiTE concentration (***Figure 3b***). The model predicted this bell-shaped relationship and revealed that high BiTE concentrations (> 100 ng/ml) would reduce the formation of ternary complexes, partly because individual antigens (CD3 and CD19) were almost completely occupied by one arm of the BiTE, limiting crosslinking with opposing cells (***Supplementary Figure S6***). IS formation increased over time and plateaued around 60 min (***Figure 3c***). We therefore restricted our incubation to 60 min considering IS quantification could be biased by serial cell-cell engagements and potential cell lysis (***Fousek et al., 2021***).

The effect of CD3 expression on IS formation was relatively small, especially at low BiTE concentrations (***Figure 3d***). The model revealed that only a small fraction of CD3 was occupied; therefore, we concluded CD3 expression was not a key driver of IS formation at low BiTE concentrations. In contrast, we found CD19 expression on target cells profoundly impacted IS formation (***Figure 3e***). These results were also predicted by the base model.

Our model predicted that cell density would also be critical to IS formation on a per-cell basis. Increasing total cell density (E+T) from 1 to 8 million per ml at an E:T ratio around 1:1 drastically boosted IS formation from 3.1% to 15.3% at 20 ng/ml BiTE and 3.9% to 27.8% at 100 ng/ml BiTE (***Figure 3f***). IS formation was further increased with high CD3-expressing effector cells at high cell densities (***Figure 3g***).

The E:T ratio also played a pivotal role in IS formation. IS formation was optimized when the E:T ratio was around 1 (***Figure 3h***). Changing the E:T ratio led to variations in the fraction of effector and target cells involved in IS formation, as predicted by the model. With higher E:T ratios, a greater fraction of target cells but lower fraction of effector cells was involved in IS formation (***Figure 3i, Supplementary Figure S10***).

Overall, we found multiple factors to be influential to IS formation. The model reasonably recapitulated IS formation dynamics under various conditions (***Figure 3b-i, Supplementary Figure S10***). The goodness of model predictions is provided in ***Supplementary Figure S11***. With good model predictability, we further investigated the influence of CD3 and CD19 binding affinities. Counterintuitively, higher affinities to CD3 resulted in lower predicted IS formation, perhaps due to higher induction of CD3 downregulation (***Supplementary Figure S12a***). Reduced IS formation at high CD3 affinities also resulted in a bell-shaped relationship (***Supplementary Figure S12b***). Notably, the approved BiTE, blinatumomab, has affinity for CD3 within the optimal range. In contrast, BiTEs with higher affinity to CD19 were predicted to enhance IS formation (***Supplementary Figure S12c and d***).

### IS variants were prevalent and well predicted by the base model

Many types of IS variants were observed in the experimental system. In total, six types of IS were quantified, including typical IS (ET), ETE, ETT, ETET, ETEE, and ETTT. IS variants with more than 4 cells were not analyzed in our study, nor included in the base model, due to their low abundance. The frequency of these variants was recorded and compared under each experimental condition.

Depending on the experimental condition, approximately 12-25% of IS observed were IS variants, although this increased up to 50% under condition 5 due to high cell density and BiTE concentration (***Figure 4a***). Among these IS variants, ETE and ETT were the most frequently observed, accounting for more than 60% of total IS variants formed under all conditions (***Figure 4b***). Conditions 5 and 11 showed high ETE frequencies due to a high E:T ratio (6:1), while conditions 6 and 12 showed high ETT frequencies due to a low E:T ratio (1:5.8). The base model well predicted the relative fraction of each IS variant under all tested conditions (***Figure 4a and b***). In general, the fraction of IS variants increased with total IS abundance. The positive correlation between the fraction of IS variants and effector cells involved in IS was well predicted by the base model (***Figure 4c***)

**Figure 4.**
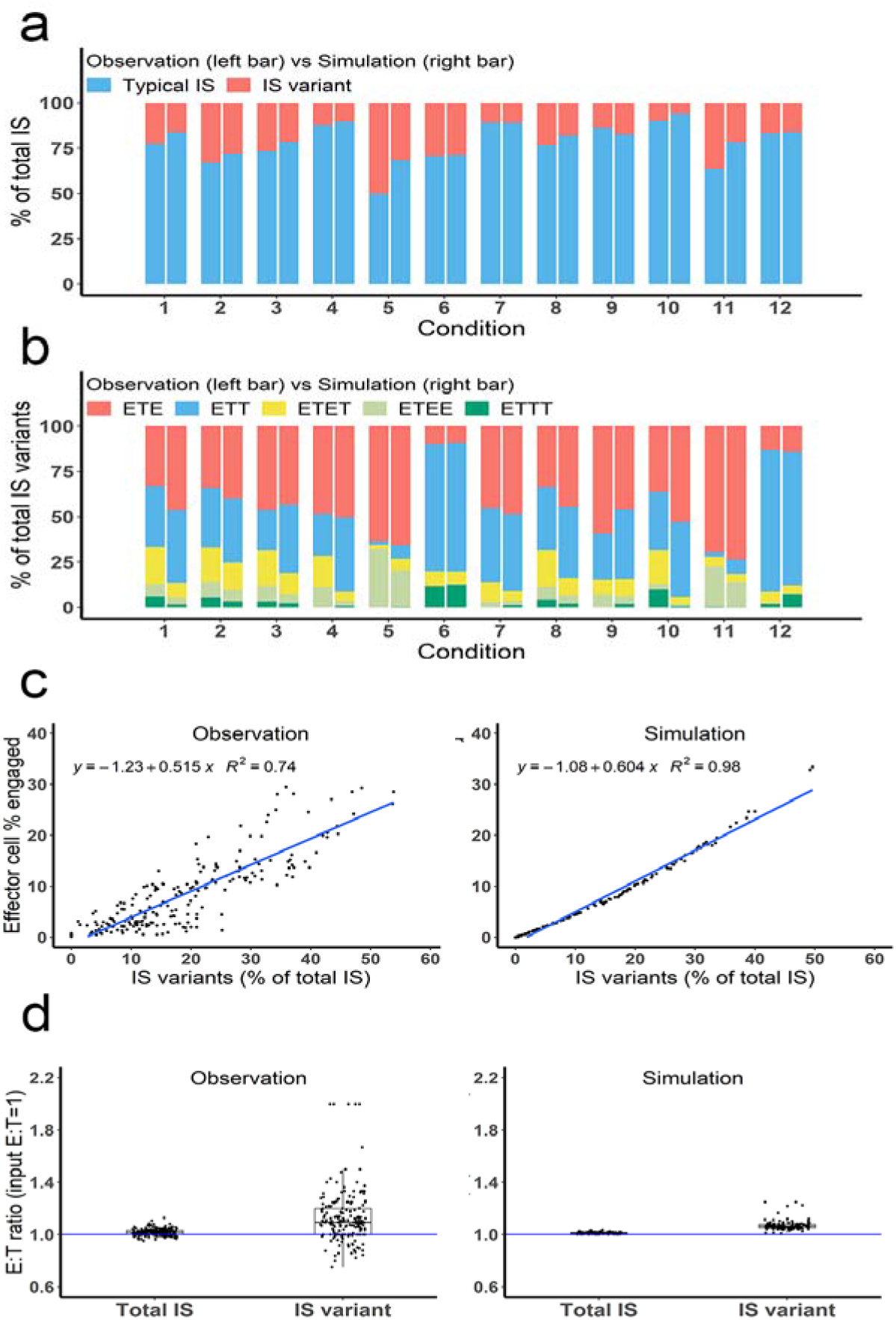
Multiple types of IS variants were observed and well predicted by the base model. In total, six types of IS were quantified, including typical IS, ETE, ETT, ETET, ETEE, and ETTT. **a**, The fraction of typical IS and variants under different conditions; **b**, The composition of IS variants (ETE, ETT, ETET, ETEE, ETTT) under different conditions; **c**, The positive correlation between the fraction of IS variants (% of total IS) and total IS formation (effector cell % engaged). The formula and R^2^ of linear regressions are shown. **d**, The E:T ratios involved in total IS and IS variants. Experimental setup: Condition 1, 2X, E:T(1), CD3(L), CD19(M), 100 ng/mL, 60 min; Condition 2, 4X, E:T(1), CD3(L), CD19(M), 100 ng/mL, 60 min; Condition 3, 2X, E:T(1), CD3(H), CD19(M), 100 ng/mL, 60 min; Condition 4, 2X, E:T(1), CD3(L), CD19(L), 100 ng/mL, 60 min; Condition 5, 4X, E:T(6), CD3(L), CD19(M), 100 ng/mL, 60 min; Condition 6, 4X, E:T(0.17), CD3(L), CD19(M), 100 ng/mL, 60 min; Conditions 7-12 are the same as Conditions 1-6, except with lower BiTE concentrations (20 ng/mL).

The E:T ratios of IS variants from all co-incubation samples were pooled for comparison (***Figure 4d***). The median E:T ratio in total IS was about 1.0. When excluding typical IS, this ratio increased to 1.1 for the remaining IS variants, suggesting slightly more effector cells were involved in IS variant formation than target cells, in line with model predictions.

### The in-vitro model predicted antigen escape and organ reservoirs

Effector T cells detach from IS and re-engage with other target cells in a process called serial cellular engagement. These effector T cells are also known as “serial killers” (***Fousek et al., 2021; Rogala et al., 2015***). We extended the base model to incorporate IS detachment and re-engagement (***Figure 5a***). The in-vitro model simulated IS formation and cellular cytotoxicity up to 72 hours. With serial engagement and killing, the fraction of target cell lysis increased considerably, even at low BiTE concentrations (***Figure 5b***).

**Figure 5.**
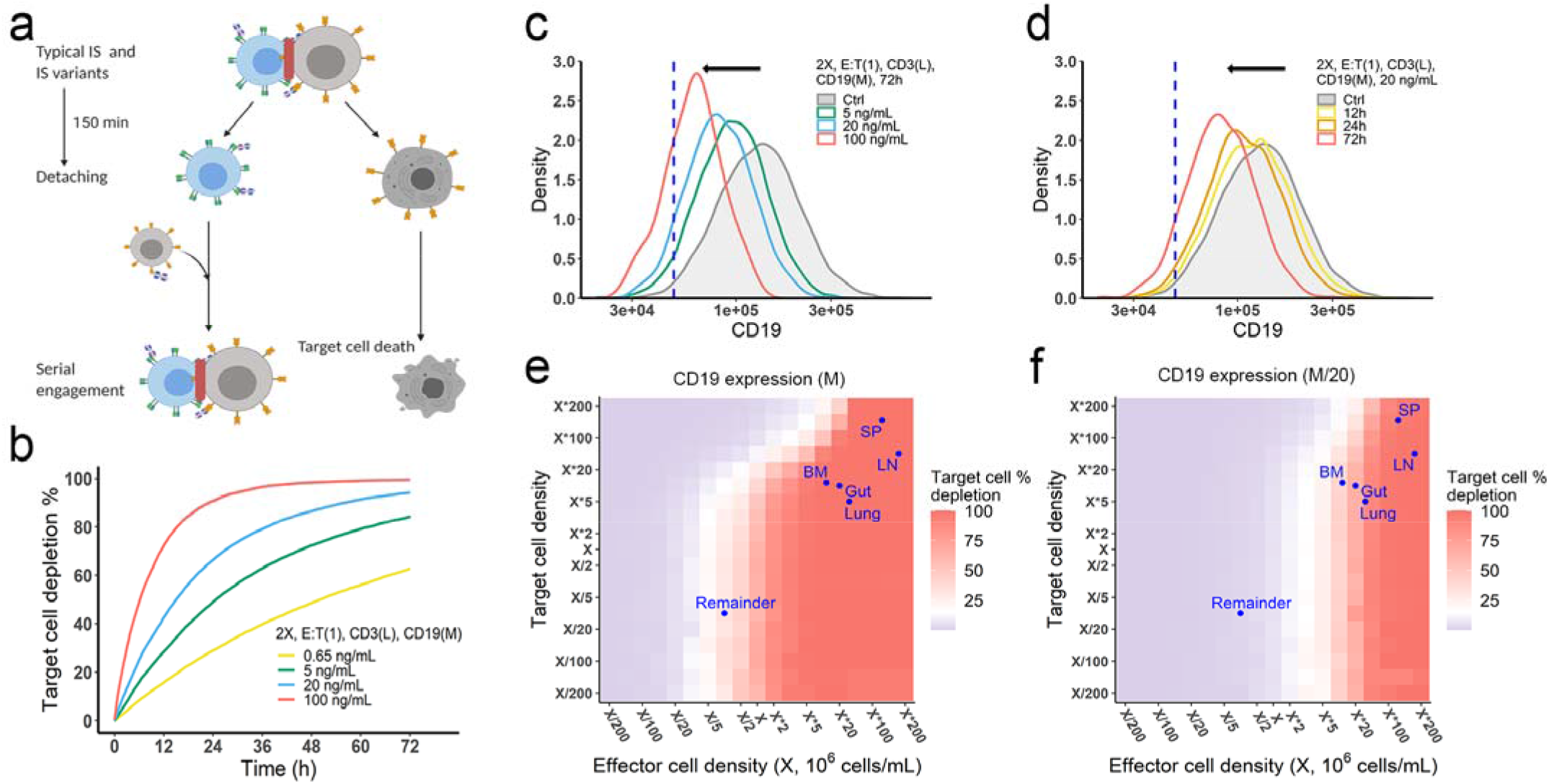
The in-vitro model predicted tumor evolution in time and space. **a**, Scheme of cell detachment and serial engagement in the in-vitro model; **b**, Long-term simulation (72h) of target cell depletion across drug concentrations; **c-d**, The effects of drug concentration (**c**) and incubation time (**d**) on CD19 expression. Dashed line, pre-defined threshold value of CD19 expression for 15% target cell depletion within 72h (initial setup: 2X, E:T(1), CD3(L), 0.65 ng/mL, 72h). Ctrl, initial distribution of CD19 expression in target cell population. **e-f**, the effects of effector and target cell density on target cell depletion (%). White, 15% target cell depletion. BM, bone marrow; LN, lymph nodes; SP, spleen; Remainder, all the rest of non-lymphoid organs. Initial setup: CD3(L), CD19(M) for (**e**), CD19 (M/20) for (**f**), 0.65 ng/mL, 72h.

Importantly, the in-vitro model predicted tumor evolution toward populations with low CD19 expression (i.e., antigen escape). Approximately 10-20% of patients who relapse after blinatumomab treatment experience antigen escape, which decreases the efficacy of subsequent anti-CD19 CAR-T cell therapy (***Braig et al., 2017; Pillai et al., 2019***). As shown in the model, tumor cells with lower CD19 expression had a lower chance of being engaged by effector cells and thus a higher probability of surviving (***Figure 5c and d***). The speed of evolution was predicted to increase at greater BiTE concentrations (***Figure 5c***) and accelerate over time (***Figure 5d***). The effect of E:T ratio, cell density, and antigen affinity on tumor evolution were also simulated (***Supplementary Figure S13***). Notably, greater IS formation led to more extensive evolution toward lower CD19-expressing cells.

The impact of cell density at clinically relevant BiTE concentrations was also interrogated (***Figure 5e and f, Supplementary Figure S14***). Notably, an increase of effector cell density resulted in higher fractions of target cell lysis at 72 hours. However, a higher density of target cells did not markedly diminish the fraction of target cells lysed at a given effector cell density, due to a compensatory increase in the probability of cell-cell encounter probability per effector cell. When target cell density was extremely high (e.g., > 5×10^6^/mL with medium CD19 expression at 1.45×10^5^/cell in ***Figure 5e***), lysis fraction decreased, as the low BiTE concentration may have become a limiting factor for IS formation. We used organ-specific effector and target cell abundance (***Supplementary Table S2***) to compare the predicted gradient of cell lysis across organs (***Figure 5e and f, Supplementary Figure S14***). Higher target cell lysis was predicted in lymph nodes and the spleen due to the abundance of effector cells in these organs. The bone marrow and all the rest of non-lymphoid organs (the remainder) showed restricted cell lysis primarily due to their relatively low abundance of effector cells (***Figure 5f***). The model also predicted that some organs like the bone marrow may become tumor cell sanctuary sites, providing space for tumor cell survival and adaptation, thereby increasing the likelihood of treatment resistance.

### The in-vivo model predicted clinical outcomes and tumor evolution across anatomical sites

We developed the in-vivo model by defining IS formation dynamics in organs and cell trafficking across organ compartments (***Figure 6a, Supplementary Methods, Supplementary Table S2 and S3***). We used the model to simulate cell lysis in each organ and tumor-killing profiles throughout the body. Organ-specific cell lysis is highly dependent on relative IS formation dynamics and thus is a function of organ-specific effector (T cell) and target (B cell) populations, as well as BiTE exposure.

**Figure 6.**
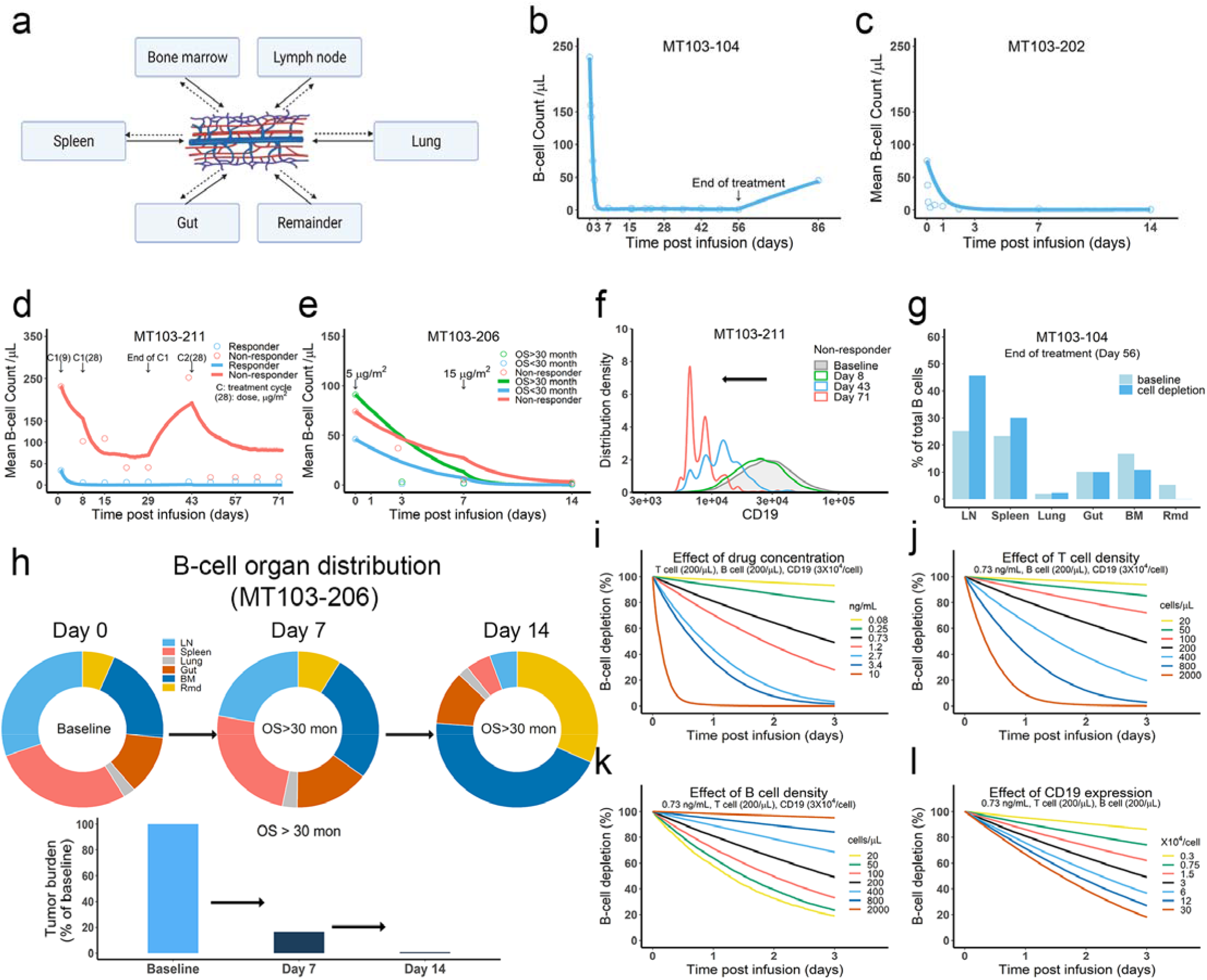
The in-vivo model predicted clinical pharmacodynamics and tumor evolution across anatomical sites. **a**, Scheme of organ compartment and cell trafficking. Remainder, all the rest of non-lymphoid organs; **b-e**, Observed and simulated patient B-cell profiles in blood. For trial information and parameters, see Supplementary Table S2 and S3; **f**, Simulated CD19 evolution in non-responder patients of trial MT103-211; **g**, Simulated cell lysis potency for each organ in trial MT103-104; **h**, Simulated baseline (Day 0), and post-treatment (Day 7 and 14) B-cell organ distribution in patients with OS > 30 month of MT103-206. Bar plot, simulated baseline and post-treatment tumor burden; **i-l**, Sensitivity analyses for the impact of drug concentration (**i**), T cell density (**j**), B cell density (**k**), and CD19 expression (**l**) on B-cell depletion. BM, bone marrow; LN, lymph nodes; OS, overall survival; Rmd, remainder.

In the in-vivo model, the blood compartment serves merely a trafficking route and does not mediate IS formation and detachment (***Figure 6a***). This assumption was supported by our observation that negligible IS was formed under shear stress forces approximating those experienced under blood flow (***Supplementary Figure S15a***). Once formed, IS in the blood are unlikely to be broken through shear stress (***Supplementary Figure S15b***). Blood B cell levels reflected the systemic average. Although only 2% of lymphocytes are present in the blood, blood flow can transport about 5×10^11^ lymphocytes each day – comparable to the total number of lymphocytes in the body (***Westermann and Pabst, 1992***).

Patients show mixed responses to BiTE immunotherapy. Some patients exhibit complete tumor eradication while others have negligible responses. By adopting patient-specific parameters, such as BiTE dosing regimens and T cell proliferation profiles, the in-vivo model reasonably predicted B-cell depletion profiles in patients (***Bargou et al., 2008; Klinger et al., 2012; Zhu et al., 2016; Zhu et al., 2018; Zugmaier et al., 2015***) treated with blinatumomab in multiple clinical trials (***Figure 6b-e, Supplementary Table S2 and S3***). In the trials MT103-211 and MT103-206, rapid accumulation of T cells in the blood was observed in responders but not non-responders (***Zhu et al., 2018; Zugmaier et al., 2015***). The model could account for these patient-specific T cell profiles and distinguish between responding and non-responding patients (***Figure 6d and e***).

Like the in-vitro model, the in-vivo model also predicted evolution toward low CD19-expressing cell populations over time, as shown in non-responders in MT103-211 (***Figure 6f***). This process is inevitable; the stronger the therapeutic pressure, the lower CD19-expression in the surviving cell population. The fraction of low CD19-expressing cells increases over time while the efficiency of tumor cell lysis decreases, leading to gradual loss of drug sensitivity.

We finally explored tumor evolution across anatomical sites and characterized the spatial gradients of cell lysis. The lymph node, spleen, and lung showed higher fractions of cell lysis than the gut, bone marrow, and remainder (***Figure 6g***). More than 45% of malignant cells in the system were lysed in the lymph nodes and around 30% were eradicated in the spleen. Lytic fractions were higher than their respective baseline levels in both organs, confirming enhanced tumor killing mediated by BiTE. In contrast, the lytic fraction in the bone marrow was lower under treatment than at baseline (***Figure 6g***), indicating poor tumor lysis efficiency. The anatomical differences in efficiency of cell lysis affected B-cell biodistribution after BiTE treatment in patients (***Figure 6h, Supplementary Figure S16***). The relative anatomical distribution of B cells also shifted considerably over time. In high responders (OS > 30 months, MT103-206), over 99% of B cells were eradicated, particularly in organs with high predicted lysis (lymph nodes and spleen). In the bone marrow, a small fraction of B cells survived that exhibited considerably lower CD19 expression than the original cell population. Unfortunately, the surviving cell populations gradually repopulated the bone marrow, leading to B-cell rebound and eventually patient relapse. By contrast, non-responders had a lower fraction of B cell lysis by day 14, with B cell distribution profiles remaining similar to the baseline (***Supplementary Figure S16***).

Sensitivity analyses confirmed that baseline tumor burden, drug concentration, cytotoxic T cell infiltration, and CD19 expression were critical to patient response (***Figure 6i-l***).

### The in-vivo model predicted optimal dosing regimens for blinatumomab

We applied the in-vivo model to simulate B cell-killing efficacy and CD19 evolution during blinatumomab treatment and compared different doses and regimens (***Figure 7a***). The initial plasma B cell abundance was assumed to be 200 cells/μL, with varying levels of growth rates. Under the approved dose (i.e., the high dose) and scheme 1, tumor-killing profiles were highly dependent upon tumor growth rate and baseline T cell abundance (***Figure 7b***). Tumors gradually accumulated resistance to treatment, especially fast-growing tumors. For slow-growing tumors with low T cell baseline, the medium dose showed comparable tumor-killing effect but resulted in less CD19 evolution than the high dose (***Figure 7c***). In contrast, for slow-growing tumors with high T cell abundance, the high dose exhibited almost complete tumor control and much less CD19 loss than the medium dose (***Figure 7d***). The high dose showed similar efficacy at two dosing schemes, but scheme 2 had fewer total doses. For fast-growing tumors, CD19 loss was significant, regardless of dose and regimen (***Figure 7e***). The medium dose at scheme 2 elicited less CD19 loss and better tumor control than scheme 1. ***Figure 7f*** summarized favorable dosing regimen under each condition. We found that the approved dose or regimen was suboptimal for almost all slow-growing tumors; rather, the medium dose or dosing regimen 2 could reach similar efficacy with slower CD19 evolution. The high dose was required for almost all fast-growing tumors, with the only exception being patients with high T cell abundance at baseline who received more benefit from the medium dose at scheme 2. Overall, our in-vivo model, through defining IS formation dynamics across anatomical sites in the system, could predict BiTE pharmacodynamics and changes in CD19 expression over time, and identify optimal dosing strategies based on baseline tumor characteristics.

**Figure 7.**
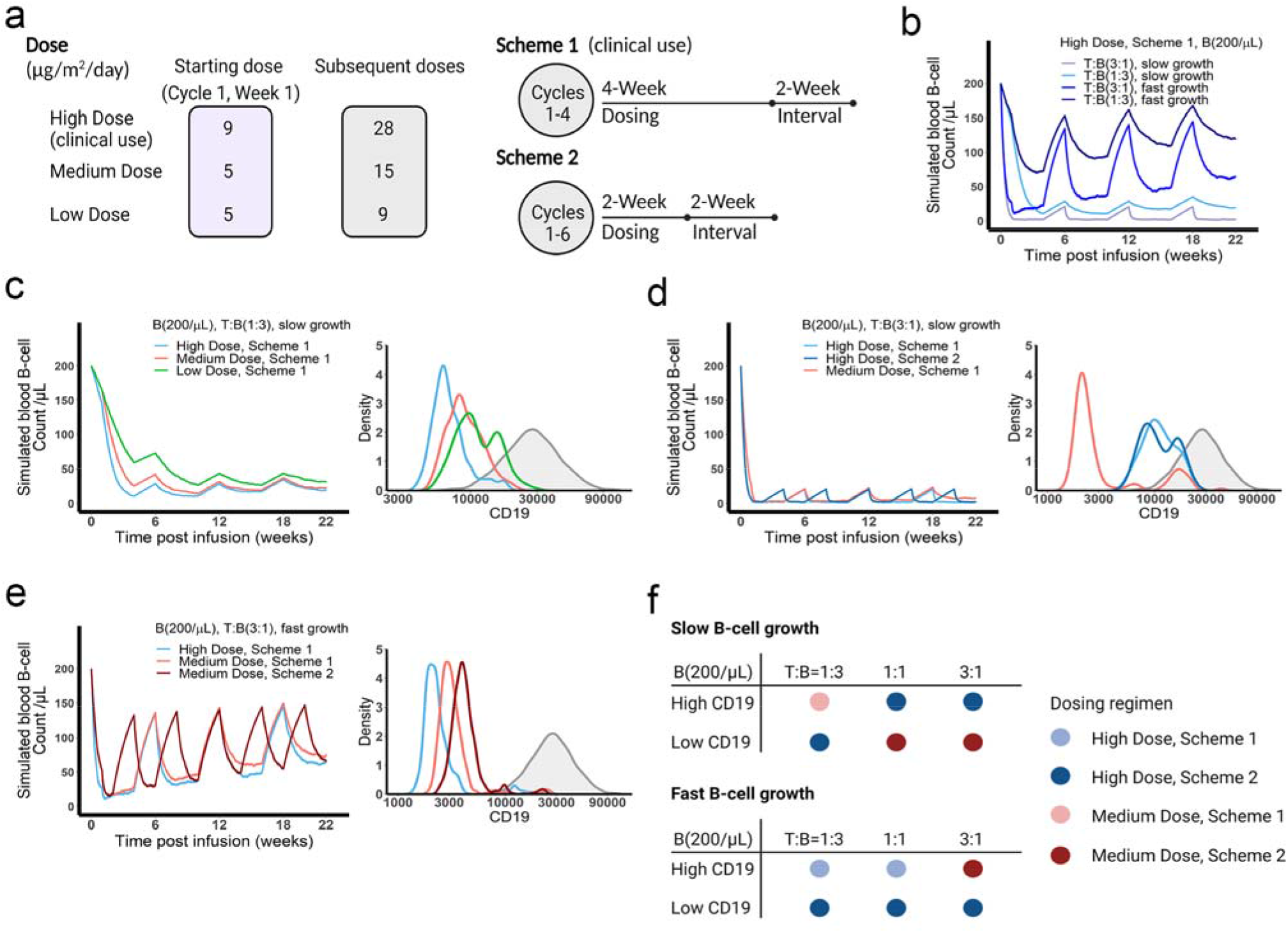
The dose and regimen of BiTE strongly influenced tumor control and CD19 evolution. **a**, Three doses (high, medium, and low) and regimens (scheme 1 and 2) were evaluated in the simulations. Starting doses were applied in the first week of cycle 1 only. The high dose with scheme 1 is the clinically approved dose and regimen of blinatumomab for the treatment of B-cell acute lymphoblastic leukemia; **b**, Simulated blood B-cell profiles under different T:B ratios and B-cell growth rates in 22-week treatment; **c-e**, Different dose levels and schemes were explored in respective conditions. Simulated blood B-cell profiles and CD19 evolution were shown. Grey line and shaded regions represent CD19 baseline expression; **f**, The favorable dosing regimen under each condition. The favorable dosing regimen was determined by comparing B-cell killing efficacy, CD19 evolution, and total dose, in this order of priority, respectively. B (200/μL), baseline B cell density in blood is assumed to 200/μL; High (**b-e**) and low CD19 expressions were the mean level of CD19 expression per B cell and set as 3×10^4^ and 1×10^4^ respectively; High growth and low growth of B cells were set to 0.071/day and 0.0071/day.

## Discussion

The clinical efficacy of BiTE immunotherapy remains suboptimal, with many of the patients who initially respond eventually experiencing disease relapse. Understanding IS formation, a crucial step in many T cell immunotherapy’s mechanism of action, can yield insights into the pharmacodynamics of BiTEs and subsequent treatment resistance. This study used an experimentally and theoretically integrated approach to examine IS formation dynamics induced by BiTEs on a population level. The abundance of IS caused by BiTEs was quantified using imaging flow cytometry, and the dynamics of IS formation were simulated with theoretical models. By defining IS formation as a spatiotemporal orchestration of molecular and cellular interactions, our theoretical models recapitulated the experimental data well. Notably, the models predicted antigen escape to be a common mechanism of resistance to BiTE immunotherapy. Tumor cells with low antigen expression accumulated over time, leading to treatment resistance and eventual disease relapse. The anatomical heterogeneity of T cell infiltration and E:T ratios across organs also conferred heterogeneous degrees of cell lysis. In particular, a subset of tumor cells in “sanctuary sites”, such as the bone marrow, may be relatively protected from effector cell lysis and fuel tumor evolution and disease relapse.

IS formation induced by BiTEs is determined mainly by two cell-scale interaction processes: cell-cell encounter and adhesion. Cell-cell encounter is the first, and in many instances, the rate limiting step to IS formation. For simplicity, the models assume random cell motion without consideration for directed or chemotactic movement (***Celli et al., 2012***). Cell density is another critical factor; cell encounter probability could become the rate-limiting step for IS formation when the target cell density is sufficiently low that effector cells have little chance of encountering target cells. This could be particularly challenging for patients with minimal residual disease. When effector cell density is low, as in the bone marrow, tumor cells have a higher chance of surviving for long enough to develop immune evasion mechanisms, leading to treatment resistance. On the other hand, when target cell density is extremely high within an organ, target cell lysis may be compromised by insufficient antibody concentrations at clinically utilized doses (***Figure 5e and f***). This is consistent with clinical observations that the efficacy of blinatumomab is much higher in patients with relatively low tumor burden (***Topp et al., 2015; Viardot and Bargou, 2018***).

The molecular crosslinking between BiTEs and antigens affects the probability of cell-cell adhesion upon encounter. Drug concentration, binding affinity, and antigen expression are critical determinants of this process. Many studies have reported a bell-shaped drug concentration-response profile for BiTE immunotherapy (***Betts et al., 2019; Douglass et al., 2013; Schropp et al., 2019; Vyver et al., 2020***), with the primary mechanism underlying the phenomenon being the oversaturation of T cell receptors at high BiTE concentrations. The theoretical model reported herein also predicts a bell-shape concentration – IS curve, but the predicted curve peaked at higher antibody concentrations if not including the possibility of CD3 down-regulation by effector cells upon antibody engagement. Our model also suggested that blinatumomab has affinity for CD3 within the optimal range of 10^−7^ – 10^−6^ M (***Supplementary Figure S12a***).

We explored the different effects of CD19 on cellular and molecular processes. Total CD19 in the system was jointly influenced by CD19 expression on membrane and target cell density. CD19 expression influenced cell lysis to a similar extent as target cell density when both factors were low (***Supplementary Figure S17a***). However, an increase of CD19 expression beyond 1.45×10^4^ receptors/cell did not further improve cell lysis. In contrast, target cell densities seemed to have a bidirectional effect on cell lysis. At low levels, escalating cell densities enhanced the probability of cell-cell encounter, while at high target cell densities, BiTE concentrations were insufficient to mediate meaningful IS formation, resulting in fewer cell-cell adhesion events and less cell lysis (***Supplementary Figure S17a and b***). This is consistent with clinical simulations (***Supplementary Figure S17c***). The different roles of CD19 expression and target cell density highlight the importance of cellular-scale interactions to IS formation that cannot be appropriately described by molecular crosslinking alone. Because of these cellular processes, our theoretical models fundamentally differ from previous BiTE pharmacodynamics models that consider molecular crosslinking only (***Betts et al., 2019; Jiang et al., 2018; Schropp et al., 2019; Song et al., 2021***). In our models, molecular crosslinking caused by BiTE, i.e., ternary complex formation, drove cell-cell adhesion events, whereas cell-cell encounters were modeled as an independent process.

Heterogeneity of CD19 antigen expression is a critical factor to BiTE-induced IS formation. Target cells with lower antigen expression had a lower probability of adhesion to T cells and thus a greater chance of survival. The theoretical models suggest that tumor evolution is an inevitable consequence to treatment, and that the stronger the therapeutic selection pressure, the more tumor cell populations evolve away from their pretreatment phenotype. Ultimately, the surviving tumor cells shift toward a low antigen expression population in a process known as antigen escape. Antigen escape is a common mechanism of resistance to T cell-based immunotherapy (***Aldoss et al., 2017; Topp et al., 2014***); however, the speed of tumor evolution toward antigen escape remains hard to predict. Through defining the formation of IS, our models show a proof of concept for predicting the trajectory of antigen escape based on baseline antigen expression.

Non-uniform tumor lysis effect across organs represents another barrier for therapy. Provided an anatomical space with few effector cells, tumor cells might use the bone marrow as a sanctuary site within which IS formation is infrequent. Insufficient selective pressure from effector cells might allow the regeneration of a newly resistant population of tumors cells that then repopulate other organs and accelerate systemic disease progression. This speculation is consistent with the clinical observation that patients under BiTE treatment often have relapses first detected in the bone marrow (***Locatelli et al., 2022***).

We used the in-vivo model to compare different doses and schedules of blinatumomab. We found that tumor baseline characteristics, including tumor growth rate, CD19 expression, and T cell abundance, greatly influenced tumor-killing pharmacodynamics, tumor evolution, and consequentially, the ideal dosing regimen. The clinically approved dose and regimen might become suboptimal for most slow-growing tumors. The medium dose or mild regimen could maintain an optimal balance between tumor-killing and evolutionary pressure (***Figure 7f***). However, it was not always the case that higher dose amounts (i.e., higher therapeutic pressure) resulted in faster tumor evolution. For slow-growing tumors with sufficient T cells (***Figure 7d and f***), the high dose with regimens 1 or 2 could cause nearly complete tumor eradication, thereby resulting negligible selection and limiting the total population size remaining for evolution. Faster and greater reductions in population size conferred by high doses might therefore reduce the chance of evolutionary rescue for slow-growing tumors.

In conclusion, our study investigated the dynamics of IS formation under various conditions mimicking the heterogeneous nature of tumor microenvironments. To our knowledge, these theoretical models are the first to quantify the entire BiTE-induced IS formation process. The models reveal trajectories of tumor evolution through antigen escape across anatomical sites and suggested dosing regimens that could confer tumor control in light of treatment-induced disease evolution. This work shows substantial implications for T cell-based immunotherapies.

## Materials and Methods

### Cell lines

Jurkat (Clone E6-1) and Raji cells were obtained from ATCC and maintained in RPMI1640 supplemented with 10–20% fetal bovine serum (FBS) and 1% penicillin-streptomycin. Cell lines were routinely tested to avoid mycoplasma contamination.

### Cell sorting and antigen expression quantification

Cell populations with high (H), medium (M) and low (L) antigen expression were sorted based on natural expression levels, without any genetic engineering. PE-anti-CD3 and PE-anti-CD19 (BD Biosciences, San Jose, CA) were used as staining antibodies and BD FACSAria II was used to perform cell sorting. Surface expression of CD3 and CD19 were quantitatively determined by Quantum™ MESF beads (Bangs laboratories, Fishers, IN) and BD LSR II flow cytometry (***Supplementary Figure S1***).

### Cell co-incubation and imaging flow cytometry

Effector cell (E, Jurkat), target cell (T, Raji), and anti-hCD19-CD3 BiTE (BioVision, Milpitas, CA) were well mixed and co-incubated in 1 mL medium at 37 °C. CD3 or CD19 expression, drug concentration, cell density, E:T ratio, and duration of co-incubation varied as initial setups. After co-incubation, effector cell, target cell, actin, and nucleus were stained by FITC-anti-CD7 (eBiosciences, San Diego, CA), PE-anti-CD20 (BD Biosciences, San Jose, CA), AF647-anti-phalloidin (Thermo-Fisher, Waltham, MA) and DAPI, respectively. Staining for surface and intracellular markers was performed as described previously (***Liu et al., 2019***). Samples were analyzed using Amnis ImageStream MKII (Luminex, Austin, TX). The frequency of IS was quantified using IDEAS (Luminex). The gating strategy is summarized in ***Supplementary Figure S2***.

### Cell co-incubation under shear stress

To mimic the shear stress in blood circulation, effector cell, target cell, and BiTE were well mixed in a circular pipe (internal diameter: 1.6 mm) and co-cultured (37 °C) at a certain flow velocity driven by a roller pump (Masterflex, Vernon Hills, IL). Flow velocities were adjusted to produce varying wall shear stresses, equivalent to those in the vein (1-6 dyn/cm^2^), artery (10-24, dyn/cm^2^) and capillary (20-40 dyn/cm^2^) (***Kamiya et al., 1984; Papaioannou and Stefanadis, 2005; Sebastian and Dittrich, 2018***). After co-culture, sample staining and analysis were the same as previously described.

**Modeling and simulation (see *Supplementary Methods*)**

## Data Availability

The data generated in this study are available upon request from the corresponding author.

## Acknowledgements

We thank Amgen Inc. for kindly sharing blinatumomab to support our pilot study.

## Author Contributions

Can Liu, Conceptualization, Formal analysis, Investigation, Methodology, Writing - original draft, Writing - review and editing; Jiawei Zhou, Investigation, Methodology, Writing - review and editing; Stephan Kudlacek, Methodology, Writing - review and editing; Timothy Qi, Writing - review and editing; Tyler Dunlap, Writing - review and editing; Yanguang Cao, Conceptualization, Formal analysis, Investigation, Methodology, Supervision, Funding acquisition, Writing - original draft, Writing - review and editing;

