## Supplementary material for "Population Dynamics of Immunological Synapse Formation Induced by Bispecific T-cell Engagers Predict Clinical Pharmacodynamics and Treatment Resistance": Suppl Materials

Supplementary Methods

Figures S1-S17

Tables S1-S3

References

**Supplementary Methods (Modeling and simulation)**

- **Base model**

Mechanistic agent-based models were developed to simulate IS formation dynamics in vitro and in vivo. We employed a sequential model-building strategy. First, we developed a model to capture IS formation dynamics within 1 hour, called the base model. The base model consisted of three modules at different dimensions: antibody-antigen binding (3D), cell-cell encounter, and cell-cell adhesion (2D) (***Figure 1 and 2***).

Antibody-antigen binding to form binary complexes (BiTE-CD3 and BiTE-CD19) was considered a 3D process among free molecules. Rapid equilibrium was assumed. At specific total concentrations of CD3 (A), CD19 (B), and BiTE (Y) in the co-incubation system, the equilibrium concentrations of free antigens ([A] and [B]) and binary complex ([AY] and [YB]) (***Supplementary* *Figure*** ***S3***) were solved according to a model with two competing binding ligands (***Yan et al., 2012***). Heterogeneous cell populations at equilibrium were generated by randomly assigning antigens and binary complexes to each individual cell (assign A and AY to effector cell, B and YB to target cell). The distribution of assigned antigens on the cells were consistent with measured CD3 and CD19 expressions in respective cell lines (***Supplementary* *Figure*** ***S1***, ***Supplementary* *Table S1***).

CD3 down-regulation on effector cells was also considered in our model ([***Lanzavecchia***](https://pubmed.ncbi.nlm.nih.gov/?term=Lanzavecchia+A&cauthor_id=9989490) ***et al., 1999;*** [***San José***](https://pubmed.ncbi.nlm.nih.gov/?term=San+Jos%C3%A9+E&cauthor_id=10714682) ***et al. 2000;*** [***Sousa***](https://pubmed.ncbi.nlm.nih.gov/?term=Sousa+J&cauthor_id=11093137) ***and Carneiro, 2000;*** [***Utzny***](https://pubmed.ncbi.nlm.nih.gov/?term=Utzny+C&cauthor_id=17012752) ***et al., 2006;*** [***Valitutti***](https://pubmed.ncbi.nlm.nih.gov/?term=Valitutti+S&cauthor_id=7753171) ***et al., 1995;*** [***Viola***](https://pubmed.ncbi.nlm.nih.gov/?term=Viola+A&cauthor_id=9362538) ***et al., 1997***). The rate of CD3 down-regulation was modeled as a function of surface binary complex abundance ([AY]’), which was described by an empirical equation that reached steady state around 1 hour incubation, in line with literature (***Supplementary* *Figure*** ***S4***). CD19 internalization was also introduced at a rate constant of 0.002/min (***Du*** ***et al., 2008;*** [***Ingle***](https://pubmed.ncbi.nlm.nih.gov/?term=Ingle+GS&cauthor_id=17991300) ***et al., 2008***). Changes in surface antigen abundance ([A]’, [AY]’, and [YB]’) for each cell was updated every 60 seconds in the model.

Encounters between effector and target cells are an essential step for IS formation. We adopted an approximate equation to determine the encounter probability of one effector cell meeting at least one target cell within a specific time, which is a function of the number of target cells in the system ([***Celli***](https://pubmed.ncbi.nlm.nih.gov/?term=Celli+S&cauthor_id=22995897) ***et al., 2012***). We assumed that effector cells diffuse independently, moving in Brownian motion, while target cells were immotile. For simplicity, the IS and IS variants were considered as a singular moving cell entity when calculating the probability of encountering an additional free cell. Notably, spatial factors were considered in this encounter probability. The equations and parameters are provided in ***Supplementary* *Figure*** ***S5***; the spatial coefficients for different encounter scenarios are listed in ***Supplementary* *Table*** ***S1***. Encounter probabilities were re-calculated every 60 seconds in the model due to the changing number of free cells over time.

When an effector cell physically contacts a target cell, it may have a chance to adhere or diffuse away. Cell-cell adhesion is mediated by ternary complexes (AYB, bond) formed from binary complexes (AY or YB) and the availability of free antigen (B or A) on opposing cell surfaces (2D binding) (***Supplementary* *Figure*** ***S6***). We assumed that stable adhesion relied on generating sufficient AYB bonds within a short contact duration. Differential equations were used to simulate the number of AYB bonds. The relationship between adhesion probability and the bond number was described by a modified deterministic equation (***Supplementary* *Figure*** ***S7;*** [***Chesla***](https://pubmed.ncbi.nlm.nih.gov/?term=Chesla+SE&cauthor_id=9726957) ***et al., 1998;*** [***Huang***](https://pubmed.ncbi.nlm.nih.gov/?term=Huang+J&cauthor_id=20357766) ***et al., 2010***). A higher number of AYB bonds yielded a higher chance for engagements to result in an IS. The randomness arose from randomly assigned antigen expressions on both cells and contact duration (0.1-5 s). It is noteworthy that the “on” and “off” rate constants used in the equations are 2D kinetic constants on cellular membrane, which were derived from 3D rate constants by a “single-step model” (***Supplementary* *Figure*** ***S8; Bell, 1978;*** [***Dreier***](https://pubmed.ncbi.nlm.nih.gov/?term=Dreier+T&cauthor_id=12209608) ***et al., 2002; Faro et al., 2017;*** [***Jansson***](https://pubmed.ncbi.nlm.nih.gov/?term=Jansson+A&cauthor_id=21044568)***, 2010***).

In the base model (time scale ≤ 1h), each free cell had only one chance to encounter in each round (60 seconds). Cells that failed to encounter or adhere would remain free cells in the next round. Newly formed IS would have a chance to encounter an additional free cell to form an IS variant in the next round (***Figure 2***). Only one free cell was allowed to be added at a time. IS variant of up to four cells were allowed in the model. The algorithm for the base model is provided in ***Supplementary* *Figure*** ***S9***. Other important assumptions in the base model include: 1) no cell proliferation and death within 1 hour; 2) no change on binding equilibrium for binary complex within 1 hour; 3) once formed, IS are not breakable.

- **In-vitro model**

Next, the in-vitro model was developed by incorporating serial cell engagement into the base model (***Figure 1b and 5a***). The duration of IS was assumed to be 150 min ([***Fousek***](https://pubmed.ncbi.nlm.nih.gov/?term=Fousek+K&cauthor_id=32205861) ***et al., 2021***), and the detached effector cells from IS became free cells for additional IS formation. We made the following updates and assumptions to extend the time scale to 72 hours: 1) binding equilibrium for binary complex was recalibrated every hour; 2) CD19 internalization was ignored; 3) CD3 down-regulation remained unchanged after 1 hour; 4) BiTE concentration remained constant.

- **Clinical translation (in-vivo model)**

Lastly, we expanded the model to develop the in-vivo model for simulating tumor-killing profiles in patients. Several additional modules, including multiple organ compartments and cell trafficking across organs, were defined in the in-vivo model (***Figure 1b and 6a***).

In the in-vivo model, IS formation occurred within each organ-specific environment (***Supplementary* *Table*** ***S2***). BiTE concentrations and antigen expression (CD3 and CD19) in human T and B cells were estimated based on reported values (***Ginaldi et al., 1996; Ginaldi et al., 1998; Haso et al., 2013; Jiang et al., 2020; Ramakrishna et al., 2019; Rosenthal et al., 2018***). The patient-specific T and B cell densities at the organ level were derived based on the blood T and B cell numbers and the predefined “partition repertoire”, which reflected the relative cell abundance between blood and each organ (***Hall et al., 2012;*** [***Hassan***](https://pubmed.ncbi.nlm.nih.gov/?term=Hassan+HT&cauthor_id=15459256) ***and*** [***El-Sheemy***](https://pubmed.ncbi.nlm.nih.gov/?term=El-Sheemy+M&cauthor_id=15459256)***,*** ***2004; Westermann and Pabst, 1992***).

The cell trafficking rate was set at 4.17% per hour, which represents the fraction of B cells trafficking through the blood during a day relative to the total B cell in the system, in line with clinical data (***Westermann and Pabst, 1992***). Each hour, 4.17% of B cells in each organ were randomly reassigned to a new location according to the pre-defined “partition repertoire” representing trafficking through the blood. T cell trafficking was not modeled, as T cell death was ignored. B cell proliferation and changes in T cell densities and BiTE dose over time were allowed (***Supplementary* *Table*** ***S2 and S3***).

As the model output, blood B cell levels after treatment were derived from the residual B cells in organs based on the pre-defined “partition repertoire”.

- **Data source and software**

Publicly available clinical data (***Bargou et al. 2008;*** [***Klinger***](https://pubmed.ncbi.nlm.nih.gov/?term=Klinger+M&cauthor_id=22592608) ***et al., 2012; Zhu et al., 2016; Zhu et al., 2018;*** [***Zugmaier***](https://pubmed.ncbi.nlm.nih.gov/?term=Zugmaier+G&cauthor_id=26480933) ***et al., 2015***) were digitized form the literature using WebPlot Digitizer. Simulation, plotting, and statistical analysis were implemented in R (3.6.0).

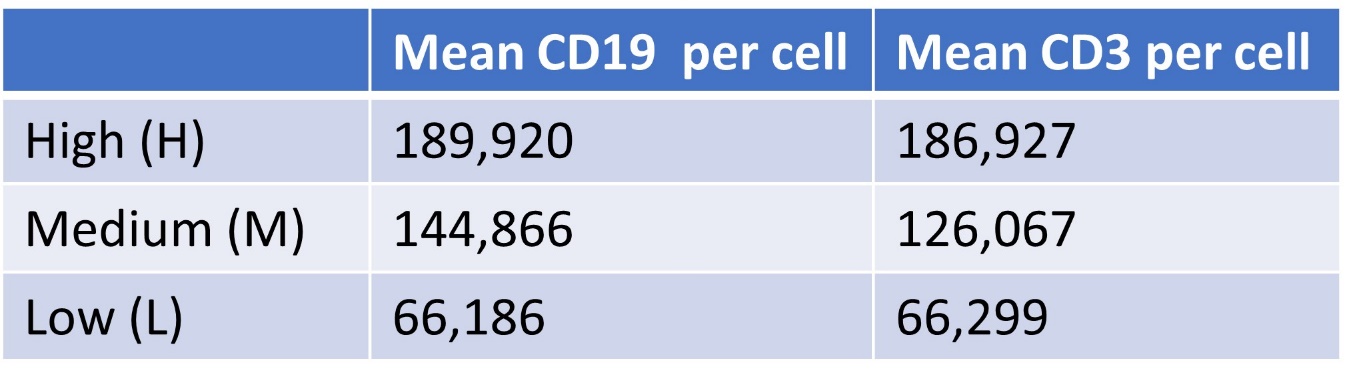

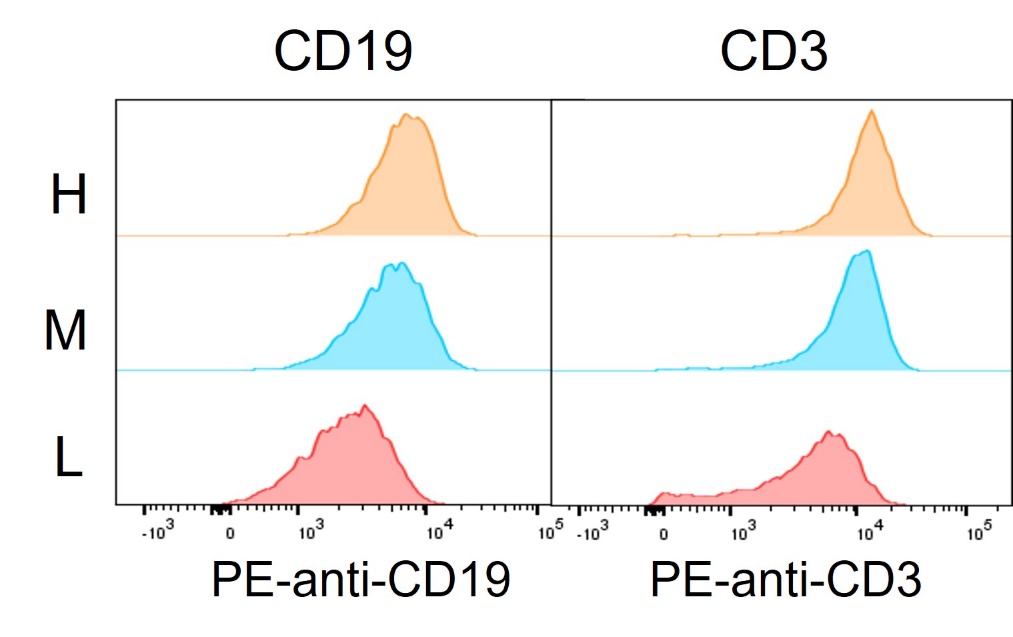

Figure S1 Histogram and quantification of high (H), medium (M) and low (L) antigen expression (CD19 and CD3) in cell subpopulations.

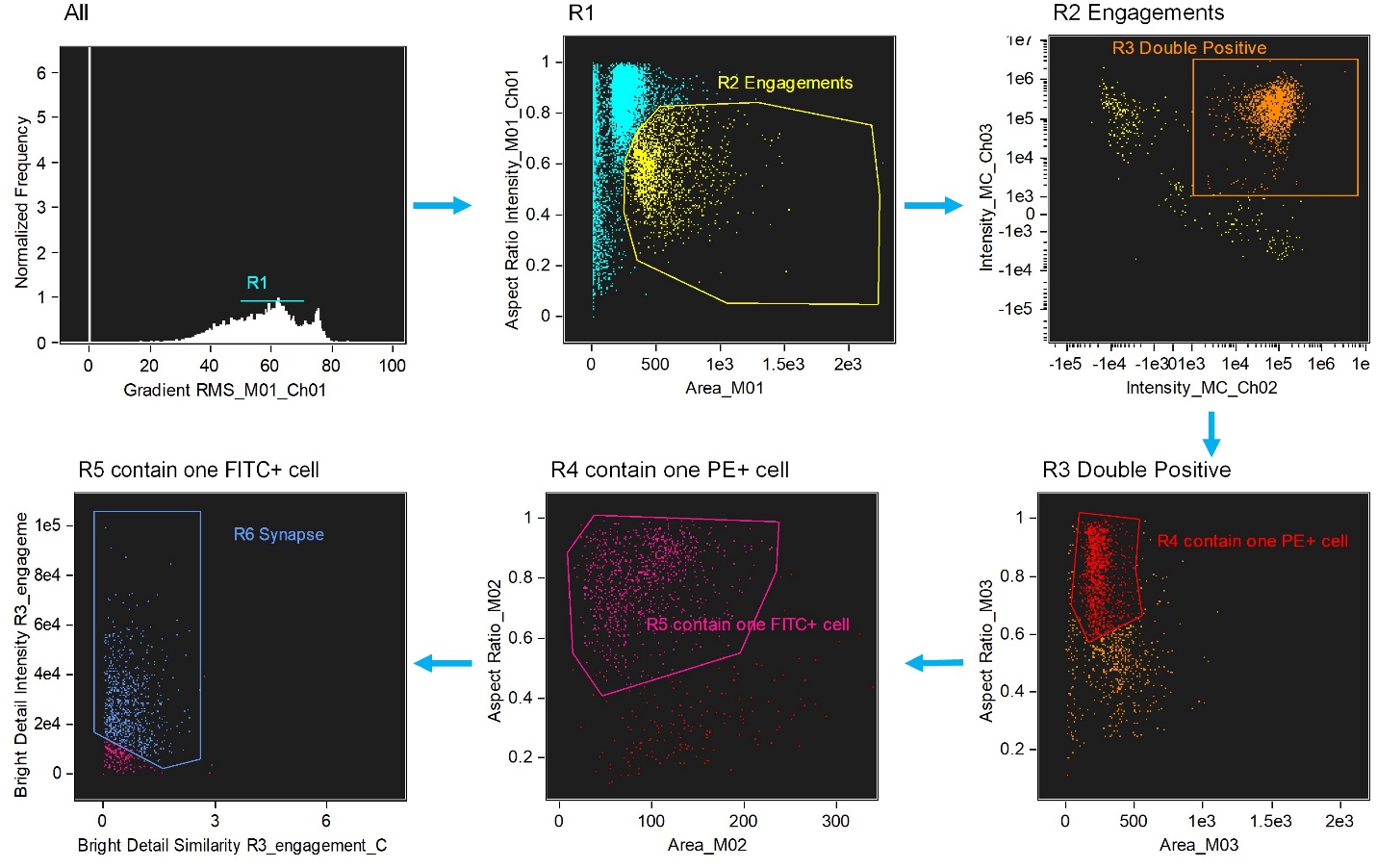

Figure S2 Image-based algorithms and gating strategy to identify typical immunological synapse (IS). Step 1, gate cells in best focus (R1). Step 2, gate conjugates (R2). Single cells (intermediate area value and high aspect ratio) are excluded. Step 3, gate CD3 and CD19 double positive (R3, FITC+PE+). Step 4, gate conjugates with only one target cell (R4). Step 5, gate conjugates with only one effector cell (R5). This subpopulation consists of doublets with only one effector and one target cell. Step 6, gate immunological synapse (R6). “Valley” mask is employed to quantify actin intensity in the region of contact (Bright detail intensity). Co-localization wizard is used to measure overlap (Bright detail similarity). For technical details, please see user’s manual of IDEAS.

**A: CD3; B: CD19; Y: BiTE**

**B**

**A**

Y

+

**YB**

**AY**

+

- Total concentrations:

${[A]}_{tot}=\left[ A \right]+\left[ AY \right]$; ${[B]}_{tot}=\left[ B \right]+[YB]$; ${[Y]}_{tot}=\left[ Y \right]+\left[ AY \right]+[YB]$	{1}

- Dissociation constants:

$K_{d, A}=\left[ A \right]\cdot\left[ Y \right]/[AY]$; $K_{d,B}=\left[ B \right]\cdot\left[ Y \right]/[YB]$	{2}

- Get binary complex concentrations ([AY] and [YB]) from equation {1} and {2}:

$[AY]=\frac{\left[ A \right]\cdot{[Y]}_{tot}/K_{d,A}}{1+\left[ A \right]/K_{d,A}+\left[ B \right]/K_{d,B}}$ ; $[YB]=\frac{\left[ B \right]\cdot{[Y]}_{tot}/K_{d,B}}{1+\left[ A \right]/K_{d,A}+\left[ B \right]/K_{d,B}}$	{3}

- Where free antigen concentrations ([A] and [B]) at equilibrium (***Yan et al., 2012***):

$[A]=\frac{{[A]}_{tot}\cdot K_{d,A}}{K_{d,A}+{[Y]}_{tot}\cdot(1-z)}$ ; $[B]=\frac{{[B]}_{tot}\cdot K_{d,B}}{K_{d,B}+{[Y]}_{tot}\cdot(1-z)}$	{4}

- Where z is the solution of a polynomial equation (for z solution see ***Yan et al., 2012***):

$If K_{d,A}\neq K_{d,B}, {the polynomial is cubic: z}^{3}+{bz}^{2}+cz+d=0$	{5}

$z satisfies, 0<z<a_{A}+a_{B} and z<1,$	{6}

$${Here, a}_{A}=\frac{\left[ A \right]_{tot}}{\left[ Y \right]_{tot}}, a_{B}=\frac{\left[ B \right]_{tot}}{\left[ Y \right]_{tot}} , k_{A}=\frac{K_{d,A}}{\left[ Y \right]_{tot}},k_{B}=\frac{K_{d,B}}{\left[ Y \right]_{tot}}$$

$b=-(2+k_{A}+k_{B}{+a}_{A}{+a}_{B})$

$c=1+2a_{A}+2a_{B}{+k}_{A}+k_{B}{+k_{B}a}_{A}{+k_{A}a}_{B}+k_{A}k_{B}$

$d=-(k_{B}a_{A}+k_{A}a_{B}{+a}_{A}{+a}_{B})$

Figure S3 *Upper*: The schema for antibody-antigen binding to form binary complex. Two antigens (CD3 (A) and CD19 (B)) competing for the same BiTE antibody (Y) was considered as a 3D binding process among free molecules, which was supposed to reach the equilibrium rapidly after co-culture initiation. *Lower*: The equations to solve free antigen ([A] and [B]) and binary complex ([AY] and [YB]) concentration levels at equilibrium (***Yan et al., 2012***).

**A: CD3; B: CD19; Y: BiTE**

- CD3 down-regulation (empirical equations):

${[AY]}_{t}^{'}={[AY]}^{'}/(1+0.1\cdot{[AY]'}^{\gamma}\cdot t^{h})$	{7}

${[A]}_{t}^{'}=\left[ A \right]^{'}/(1+0.1\cdot[{AY]'}^{\gamma}\cdot t^{h})$	{8}

where, for each effector cell, [AY]’ and [A]’ are assigned AY and A concentration on cell surface; [AY]’_t_ and [A]’_t_ are AY and A concentration on cell surface at time t; hill function: γ = 0.9; h = 0.7.

- CD19 internalization:

${[YB]}_{t}^{'}={[YB]}^{'}\cdot e^{-k_{int}t}$	{9}

where, for each target cell, [YB]’ is assigned YB concentration on cell surface; [YB]’_t_ is the YB concentration on cell surface at time t; k_int_ stands for internalization rate constant (0.002 min^-1^) (***Du*** ***et al., 2008;*** [***Ingle***](https://pubmed.ncbi.nlm.nih.gov/?term=Ingle+GS&cauthor_id=17991300) ***et al., 2008***)

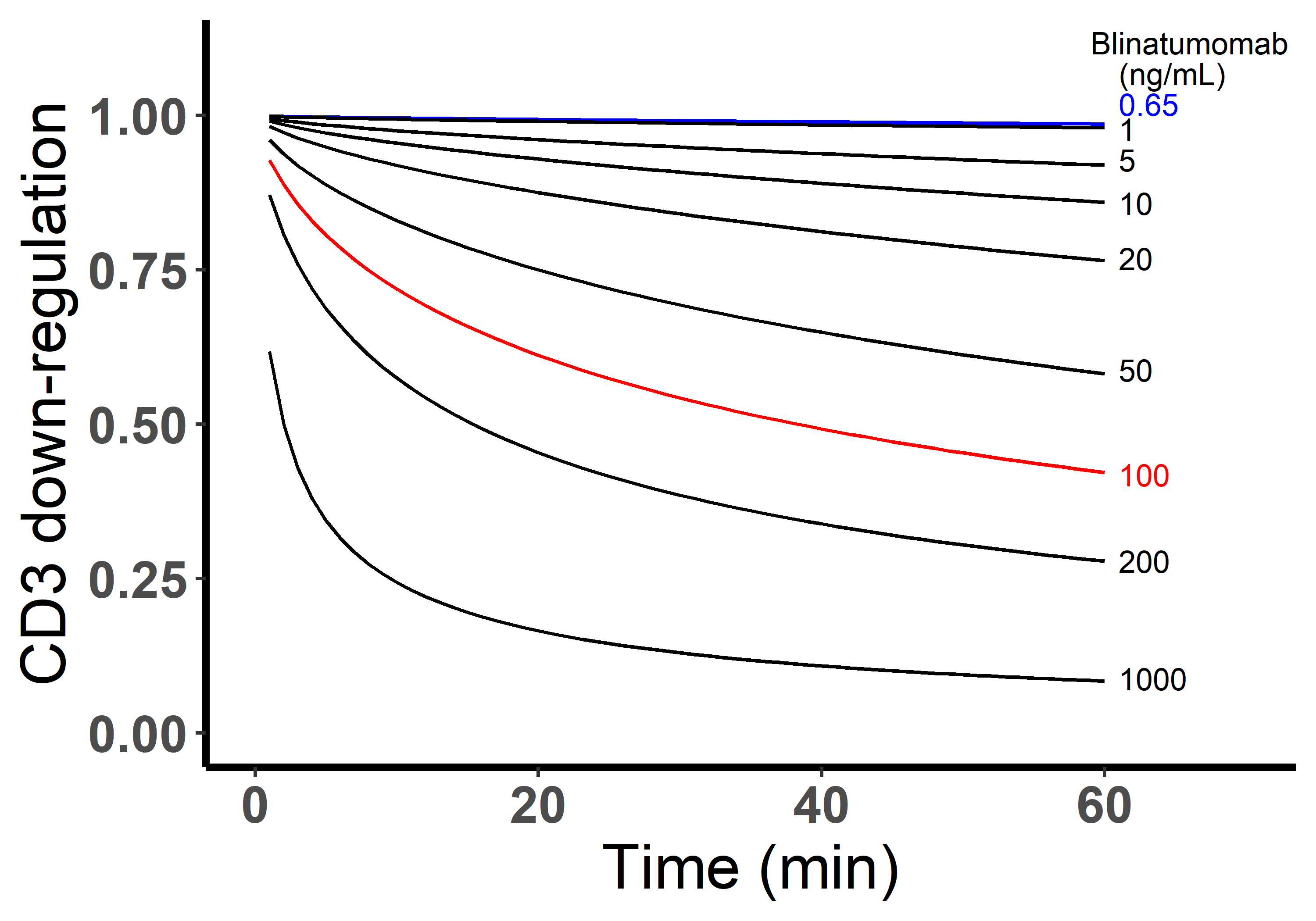

Figure S4 *Upper*: empirical equations for CD3 down-regulation and CD19 internalization within 1h after binary-complex equilibrium. *Lower*: simulated CD3 down-regulation over time at different blinatumomab concentrations. Initial setup: 2×10^6^ cells/mL, CD3 expression (L), CD19 expression (M), E:T ratio=1.

- Encounter probability ([***Celli***](https://pubmed.ncbi.nlm.nih.gov/?term=Celli+S&cauthor_id=22995897) ***et al., 2012***):

${for a single E encounter free T, P}_{e}=1-e^{-\alpha Tt}$	{10}

${for a ET encounter free E, P}_{eE}=(1-e^{-\alpha Et})\cdot2\cdot f_{ETE}$	{11}

${for a ET encounter free T, P}_{eT}=(1-e^{-\alpha Tt})\cdot2\cdot f_{ETT}$	{12}

${for a ETE encounter free E, P}_{eEE}=(1-e^{-\alpha Et})\cdot3\cdot f_{ETEE}$	{13}

${for a ETE encounter free T, P}_{eET}=(1-e^{-\alpha Tt})\cdot3\cdot f_{ETET}$	{14}

${for a ETT encounter free E, P}_{eTE}=(1-e^{-\alpha Et})\cdot3\cdot f_{ETTE}$	{15}

${for a ETT encounter free T, P}_{eTT}=(1-e^{-\alpha Tt})\cdot3\cdot f_{ETTT}$	{16}

$The \alpha is given by, \frac{1}{\alpha}=\frac{R^{3}}{3Db}-\frac{3R^{2}}{5D}$	{17}

Where, Pe, encounter probability; E and T, cell number of free effector and target cells; t, time; f, spatial coefficients, the values were listed in ***Supplementary Table S1***; α, mean hitting rate; D, cell diffusivity in solution, 0.83 µm^2^·s^-^1 (***Miller et al. 2003***); b, distance between cell centers at encounter, 11 µm (radius: effector cell, 5µm _;_ target cell, 6 µm); R, radius of spherical co-culture system, 6200 µm (1 mL system).

f_ETE_ = 0.75

f_ETT_ = 0.66

**E**

**E**

**E**

**E**

**T**

**T**

**T**

**T**

**E**

Figure S5 *Upper*: equations and parameters for encounter probability; *Lower*: spatial factor when a typical IS (ET) encounters an additional free target cell or effector cell. Based on imaging data, it was allowed a single effector cell to maximumly have 3 binding spots for target cells, whereas up to 4 effector cells for a single target cell. Dashed circles represent available spots for an additional effector or target cell binding to a typical IS (solid circles).

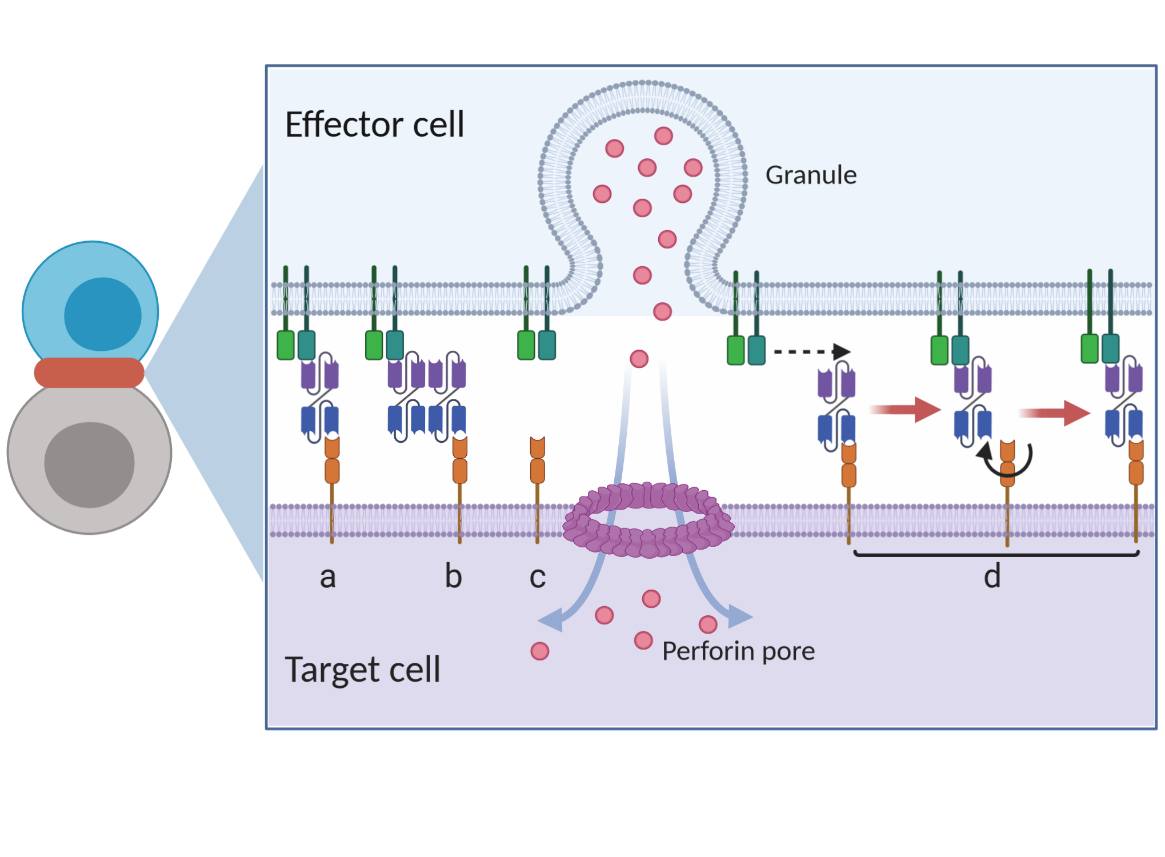

Figure S6 Graphical presentation of ternary complex (CD3-BiTE-CD19, bond) formation during cell-cell adhesion. The bond is formed by binding between a binary complex and a free antigen (**a**), rather than two binary complexes (**b**) or two free antigens (**c**). Three steps of 2-D binding on membrane (**d**): diffusion, rotation, and molecular binding.

**A: CD3; B: CD19; Y: BiTE**

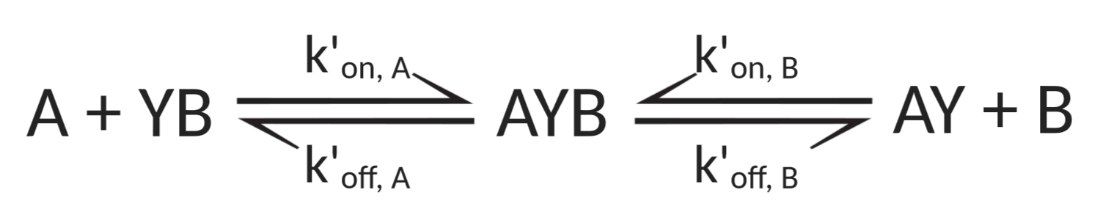

- The formation of ternary complex (AYB, bond) can be characterized as

$\frac{d{[AYB]}^{'}}{dt}=k_{on, A}^{'}\cdot\left[ A \right]^{'}\cdot\left[ YB \right]^{'}+k_{on, B}^{'}\cdot\left[ B \right]^{'}\cdot\left[ AY \right]^{'}-k_{off, A}^{'}\cdot\left[ AYB \right]^{'}-k_{off, B}^{'}\cdot\left[ AYB \right]^{'}$	{18}

$\frac{d{[A]}^{'}}{dt}=k_{off, A}^{'}\cdot{[AYB]}^{'}-k_{on, A}^{'}\cdot{[A]}^{'}\cdot{[YB]}^{'}$	{19}

$\frac{d{[AY]}^{'}}{dt}=k_{off, B}^{'}\cdot{[AYB]}^{'}-k_{on, B}^{'}\cdot{[B]}^{'}\cdot{[AY]}^{'}$	{20}

$\frac{d{[B]}^{'}}{dt}=k_{off, B}^{'}\cdot\left[ AYB \right]^{'}-k_{on, B}^{'}\cdot\left[ B \right]^{'}\cdot\left[ AY \right]^{'}$	{21}

$\frac{d{[YB]}^{'}}{dt}=k_{off, A}^{'}\cdot{[AYB]}^{'}-k_{on, A}^{'}\cdot{[A]}^{'}\cdot{[YB]}^{'}$	{22}

- Adhesion probability ([***Chesla***](https://pubmed.ncbi.nlm.nih.gov/?term=Chesla+SE&cauthor_id=9726957) ***et al., 1998;*** [***Huang***](https://pubmed.ncbi.nlm.nih.gov/?term=Huang+J&cauthor_id=20357766) ***et al., 2010***)

$N_{bond, \tau}={[AYB]}_{\tau}^{'}\cdot S_{contact}$	{23}

$Pa=1-e^{-\beta\cdot N_{bond.\tau}}$	{24}

Where, $k_{on}^{'}, k_{off}^{'}$, effective rate constants for 2-D binding on membrane, see ***Supplementary*** ***Figure S8*** and ***Supplementary*** ***Table S1;***

${[A]}^{'}, {[B]}^{'}, {[AY]}^{'}, {{[YB]}^{'}, [AYB]}^{'}$, antigen density on cell surface;

$N_{bond,\tau}$_,_ bond number at time τ;

${[AYB]}_{\tau}^{'}$, bond density at contact duration τ (randomly assigned from 0.1-5s);

$S_{contact}$, apparent contact area, ~ 5 µm^2^;

$Pa$, adhesion probability;

β, sensitive factor, 0.033 (for base and in-vitro model).

Figure S7 Equations and parameters for ternary complex formation and adhesion probability

**A: CD3; B: CD19; Y: BiTE**
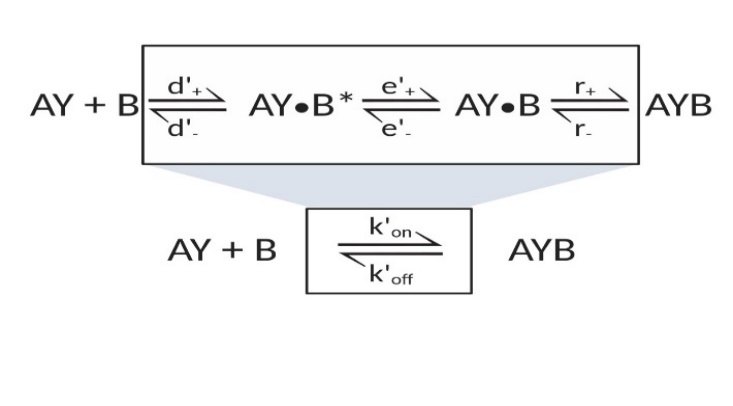

- 3-D binding in solution (***Bell, 1978; Faro et al., 2017***)

$d_{+}=4\pi\cdot R_{AYB}\cdot(D_{AY}+D_{B})$; $d_{-}=3\cdot R_{AYB}^{-2}\cdot(D_{AY}+D_{B})$	{25}

$e_{+}=0.04\cdot e_{-}$ ; $e_{-}=3\cdot d_{-}/4\pi^{2}$	{26}

; {27}

$k_{off}=\frac{d_{-}\cdot e_{-}\cdot r_{-}}{d_{-}\cdot e_{-}+r_{+}(d_{-}+e_{+})}$

$k_{on}=\frac{d_{+}\cdot e_{+}\cdot r_{+}}{d_{-}\cdot e_{-}+r_{+}(d_{-}+e_{+})}$

Similar equations were applied for A+BY$\rightleftharpoons$AYB as well

- 2-D binding on membrane (***Bell, 1978; Faro et al., 2017***)

$d_{+}^{'}=2\pi\cdot(D_{AY}^{'}+D_{B}^{'})$ ; $d_{-}^{'}=2\cdot R_{AYB}^{-2}\cdot(D_{AY}^{'}+D_{B}^{'})$	{28}

$e_{+}^{'}=0.04\cdot e_{-}^{'}$ ; $e_{-}^{'}=3\cdot d_{-}^{'}/4\pi^{2}$	{29}

$k_{on}^{'}=\frac{d_{+}^{'}\cdot e_{+}^{'}\cdot r_{+}}{d_{-}^{'}\cdot e_{-}^{'}+r_{+}(d_{-}^{'}+e_{+}^{'})}$	; $k_{off}^{'}=\frac{d_{-}^{'}\cdot e_{-}^{'}\cdot r_{-}}{d_{-}^{'}\cdot e_{-}^{'}+r_{+}(d_{-}^{'}+e_{+}^{'})}$	{30}

Similar equations were applied for A+BY$\rightleftharpoons$AYB as well

Where, $d_{+}, d_{-}$, $d_{+}^{'},$ $d_{-}^{'}$, forward and reverse rate constants of antigen encounter;

$e_{+}$, $e_{-}$, $e_{+}^{'},$ $e_{-}^{'}$, forward and reverse rate constants of antigen rotation;

$r_{+}$, $r_{-}$, chemical association and dissociation rate constants;

$k_{on}, k_{off},k_{on}^{'}, k_{off}^{'}$, effective rate constants, see ***Supplementary*** ***Table S1***;

$D_{AY},$ $D_{B}$, diffusion constant in solution, $D_{AY}=D_{B}=D_{Y}$, 50 µm^2^/s (***Bell, 1978***);

$D_{AY}^{'},D_{B}^{'}$, diffusion constant on membrane, $D_{AY}^{'}+D_{B}^{'}$= 0.01 µm^2^/s (***Bell, 1978***);

$R_{AYB}$, distance of reactants (AY and B), 5 nm ([***Jansson***](https://pubmed.ncbi.nlm.nih.gov/?term=Jansson+A&cauthor_id=21044568)***, 2010***).

Figure S8 *Upper*: “single-step model” for 2-D binding on cell membrane; *Lower*: equations for effective kinetic constants for 2-D binding on cell membrane.

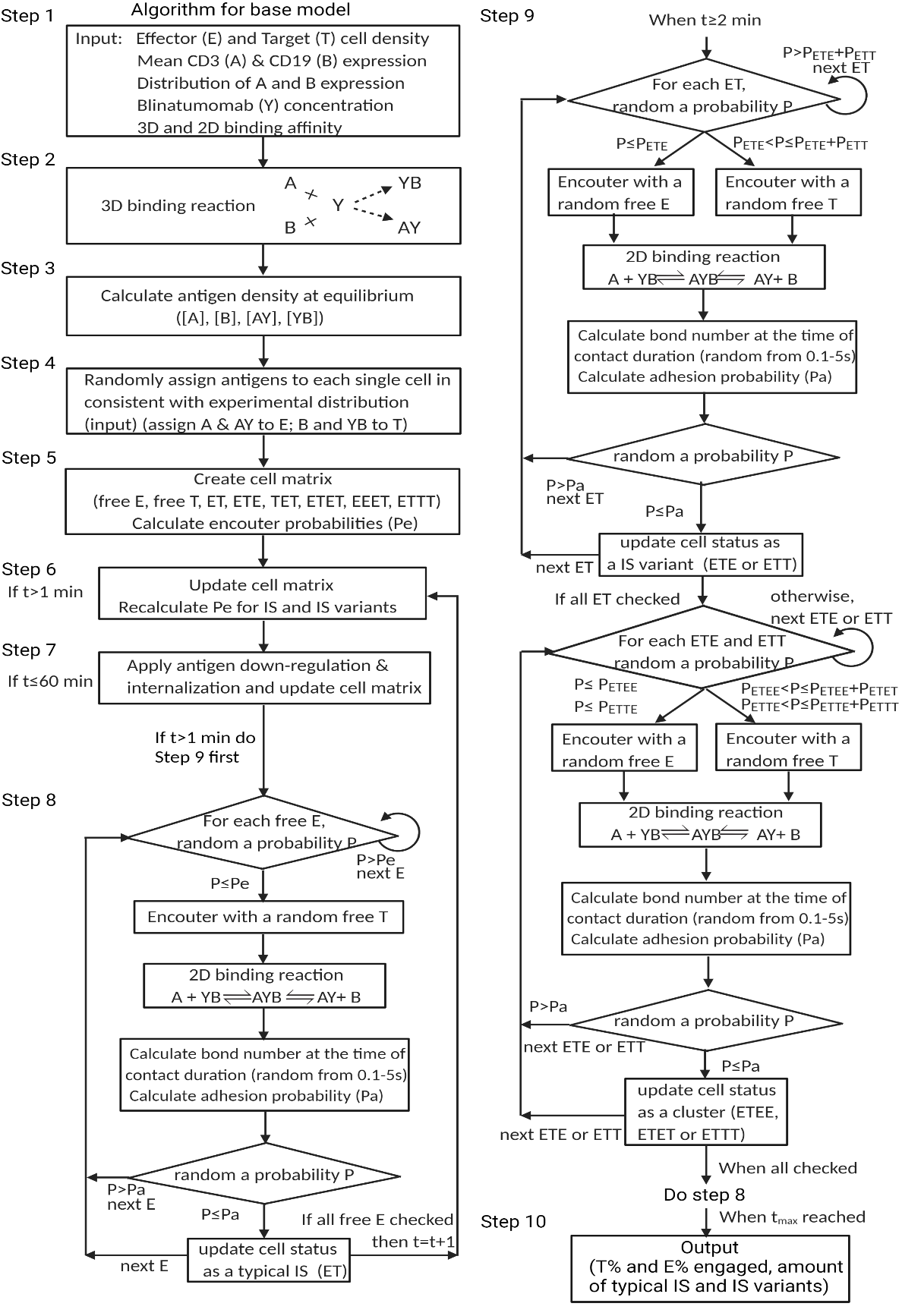

Figure S9 Algorithm for the base model

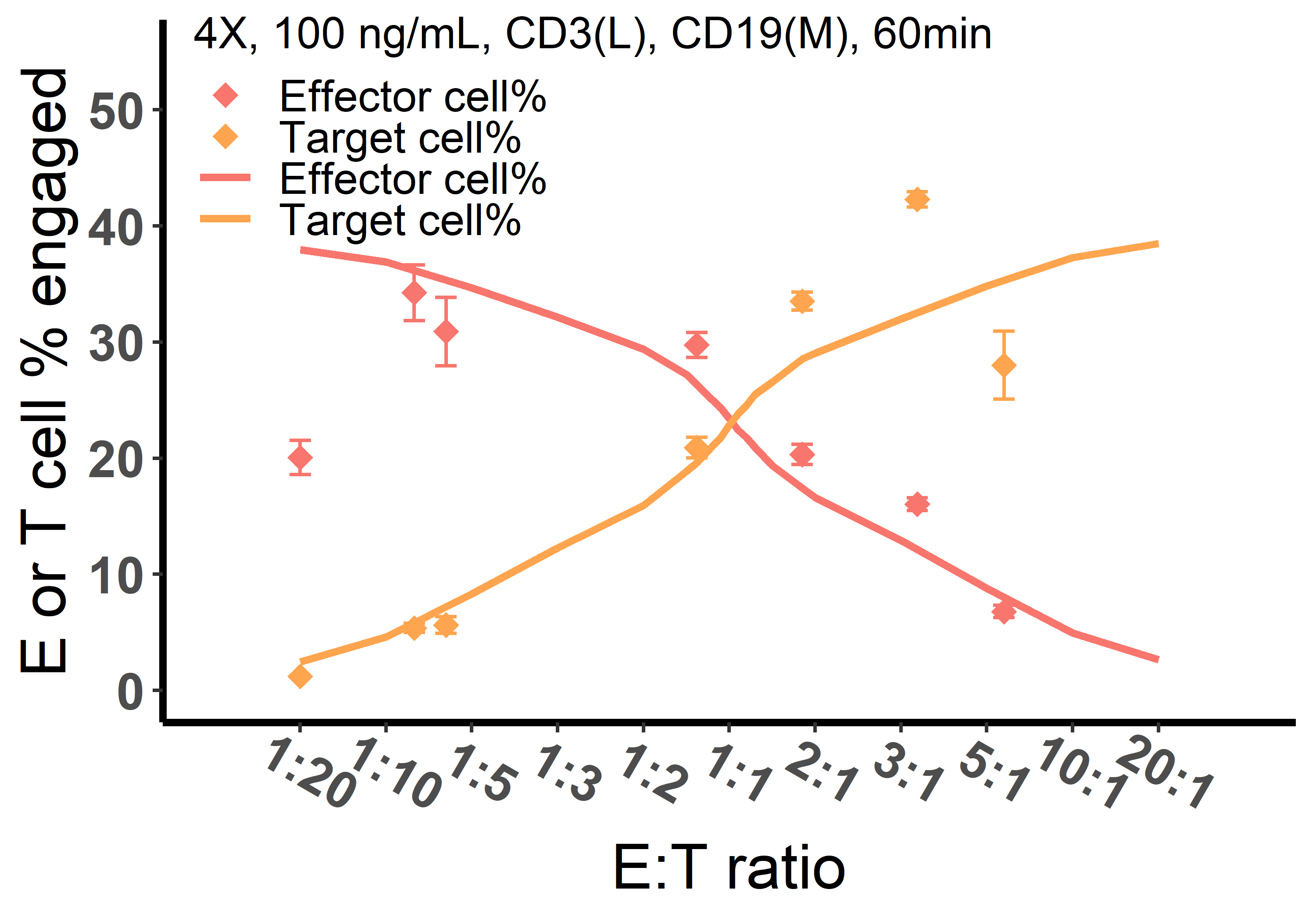

Figure S10 The effect of E:T ratio on cell-cell engagement at 100 ng/mL BiTE concentration. Red, effector cell % engaged; Orange, target cell % engaged. Observations were displayed in dots and simulations were solid curves. The base model was applied to perform simulations. Initial setup: 4X, 4×10^6^ cells/mL; 100 ng/mL, blinatumomab concentration; CD3(L), CD3 expression (Low); CD19(M), CD19 expression (Medium); 60 min, incubation duration.

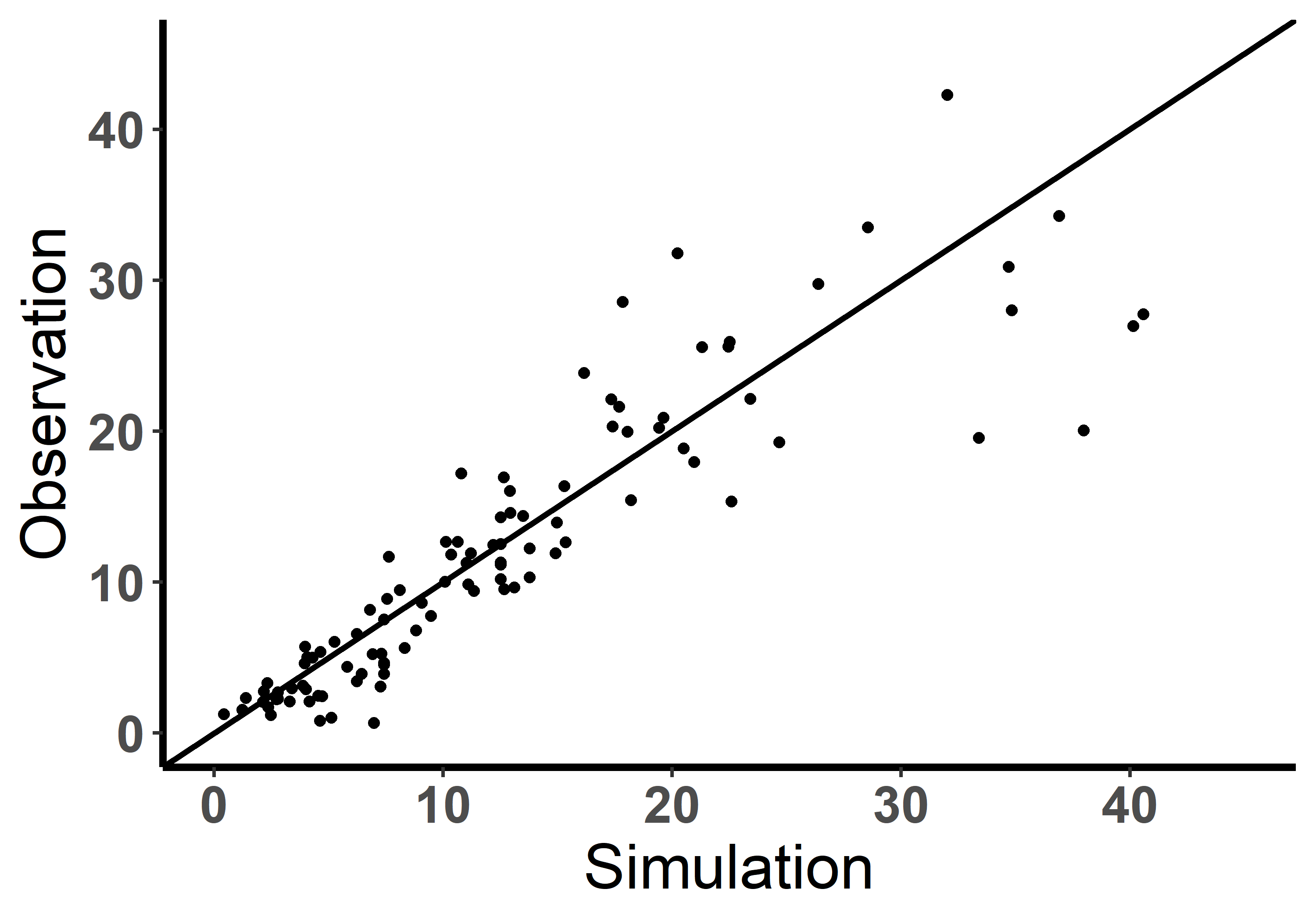

Figure S11 Performance of the base model (observation vs simulation). solid line, y = x.

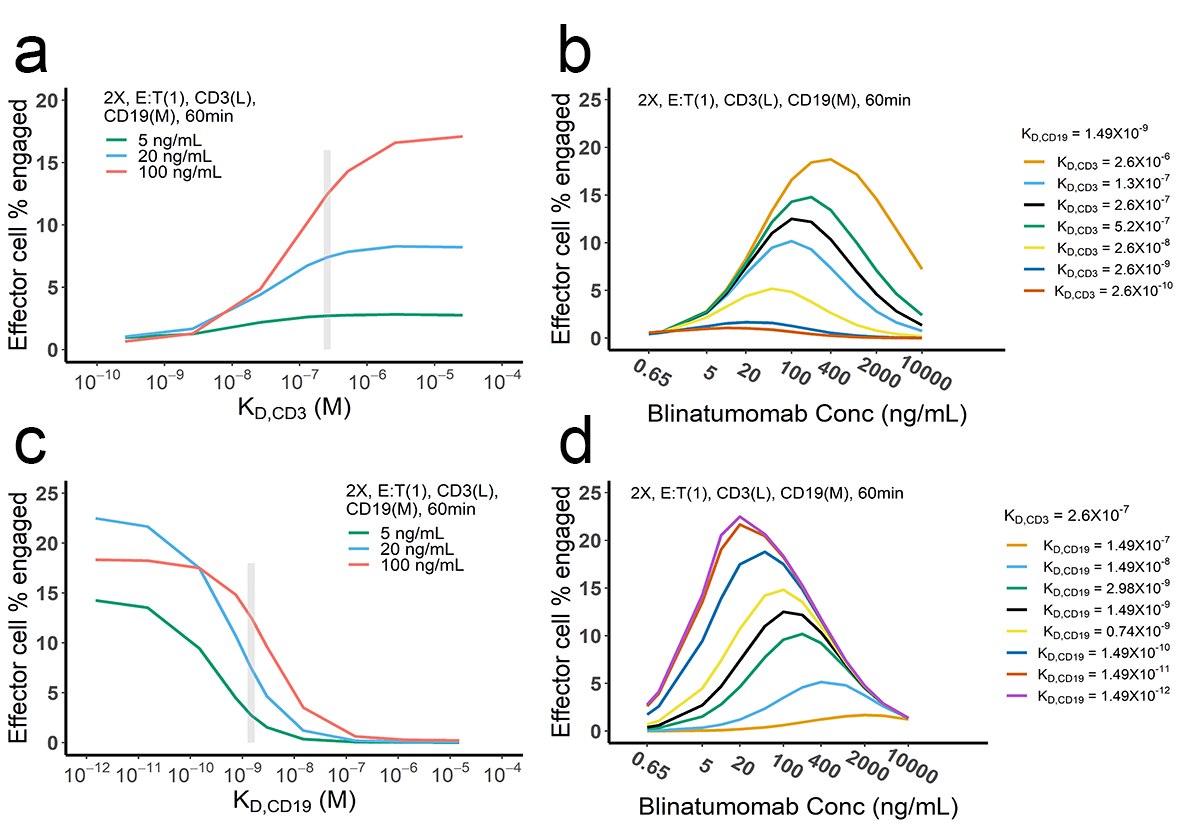

Figure S12 The effect of binding affinity on BiTE-mediated cell-cell engagement. The base model was applied to perform simulations. K_D_, dissociation constant; grey vertical line, the K_D_ value of blinatumomab;. 2X, 2×10^6^ cells/mL; E:T(1), E:T ratio = 1; CD3(L), CD3 expression (Low); CD19(M), CD19 expression (medium); 5, 20, 100 ng/mL, blinatumomab concentration; 60 min, incubation duration.

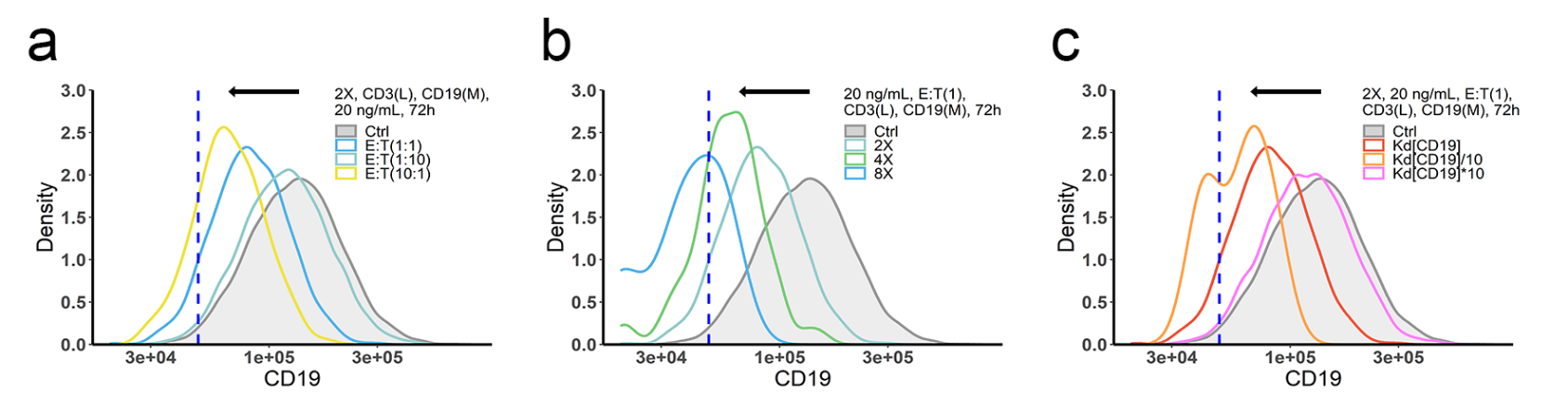

Figure S13 The effects of ET ratio (a), cell density (b) and binding affinity (c) on CD19 evolution. The in-vitro model was applied to perform simulations. Dashed line, threshold value of CD19 expression for 15% target cell depletion within 72h (initial setup: 2X, E:T(1), CD3(L), 0.65 ng/mL, 72h). Ctrl, initial CD19 distribution in target cell population. K_d_, dissociation constant; 2X, 2×10^6^ cells/mL; E:T(1), E:T ratio=1; CD3(L), CD3 expression (Low); CD19(M), CD19 expression (medium); 0.65, 20 ng/mL, blinatumomab concentration; 72h, incubation duration.

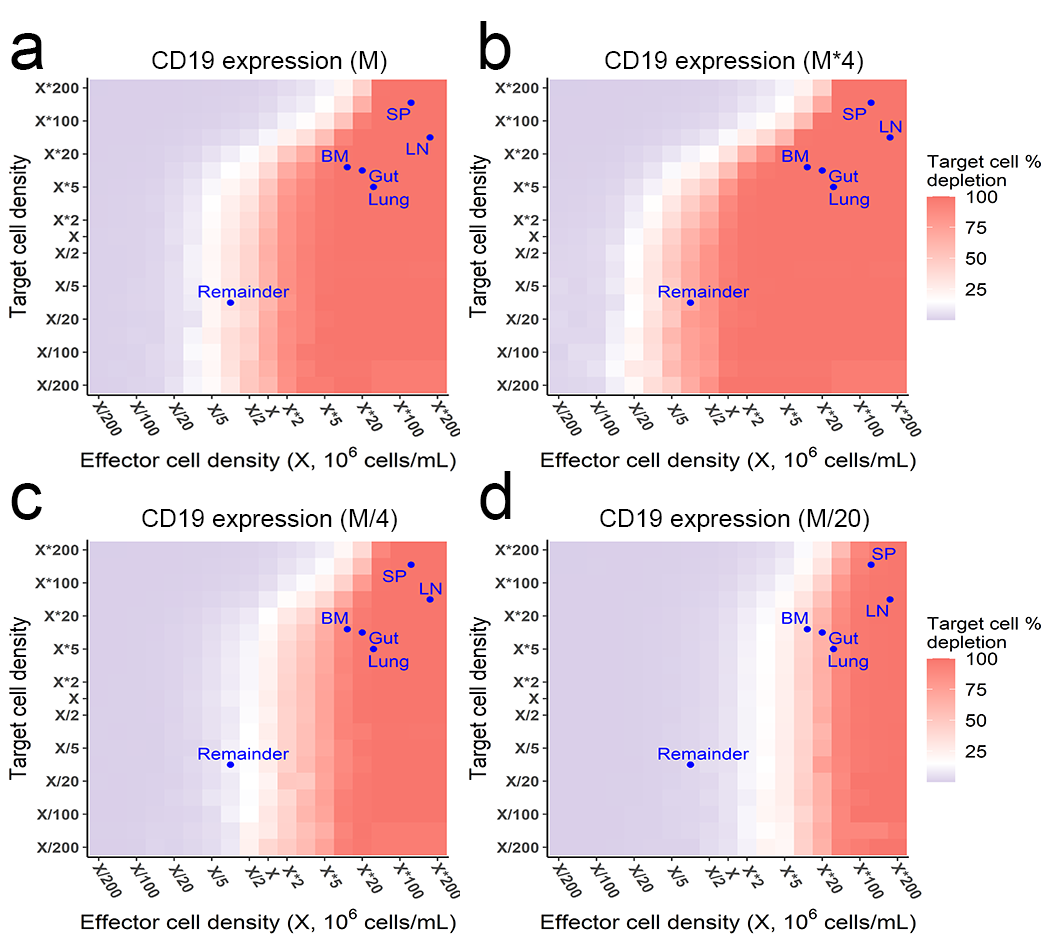

Figure S14 The effects of effector and target cell density on target cell depletion (%) at 72h. Simulations were performed by the in-vitro model. Dots indicated the effect and target cell densities in healthy human organs. White color, 15% target cell depletion. BM, bone marrow; LN, lymph nodes; SP, spleen; Remainder, all the rest of non-lymphoid organs. Initial setup: CD3(L), CD19(M) for a, CD19 (M×4) for b, CD19 (M/4) for c, CD19 (M/20) for d, 0.65 ng/mL, 72h.

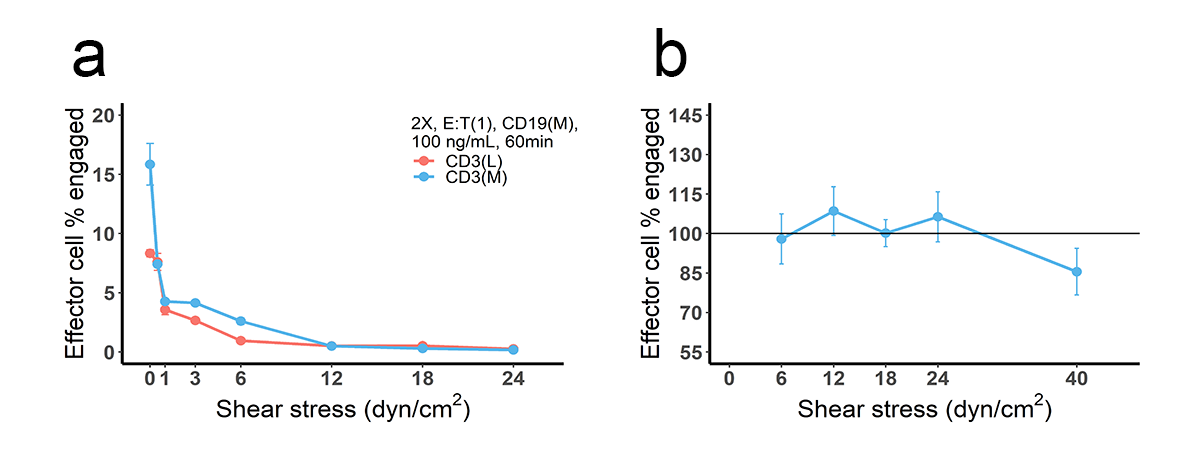

Figure S15 The effect of blood flow on production and stability of IS.

**a,** cell co-culture was conducted at different flow velocities, producing different shear stresses to mimic the effect of blood flow. Initial setup: 2X, 2×10^6^ cells/mL; E:T(1), E:T ratio=1; CD3(L), CD3 expression (Low); CD3(M), CD3 expression (medium); CD19(M), CD19 expression (medium); 100 ng/mL, blinatumomab concentration; 60 min, incubation duration.

**b,** cell incubation was conducted at static condition (initial setup: 2X, E:T(1), CD19(M), CD3(L), 100 ng/mL, 60 min, shear stress=0) and followed by adding different shear stresses (5 min) to test the stability of pre-existing IS. The reference line (100%) indicated the frequency of effector cell engaged at static condition.

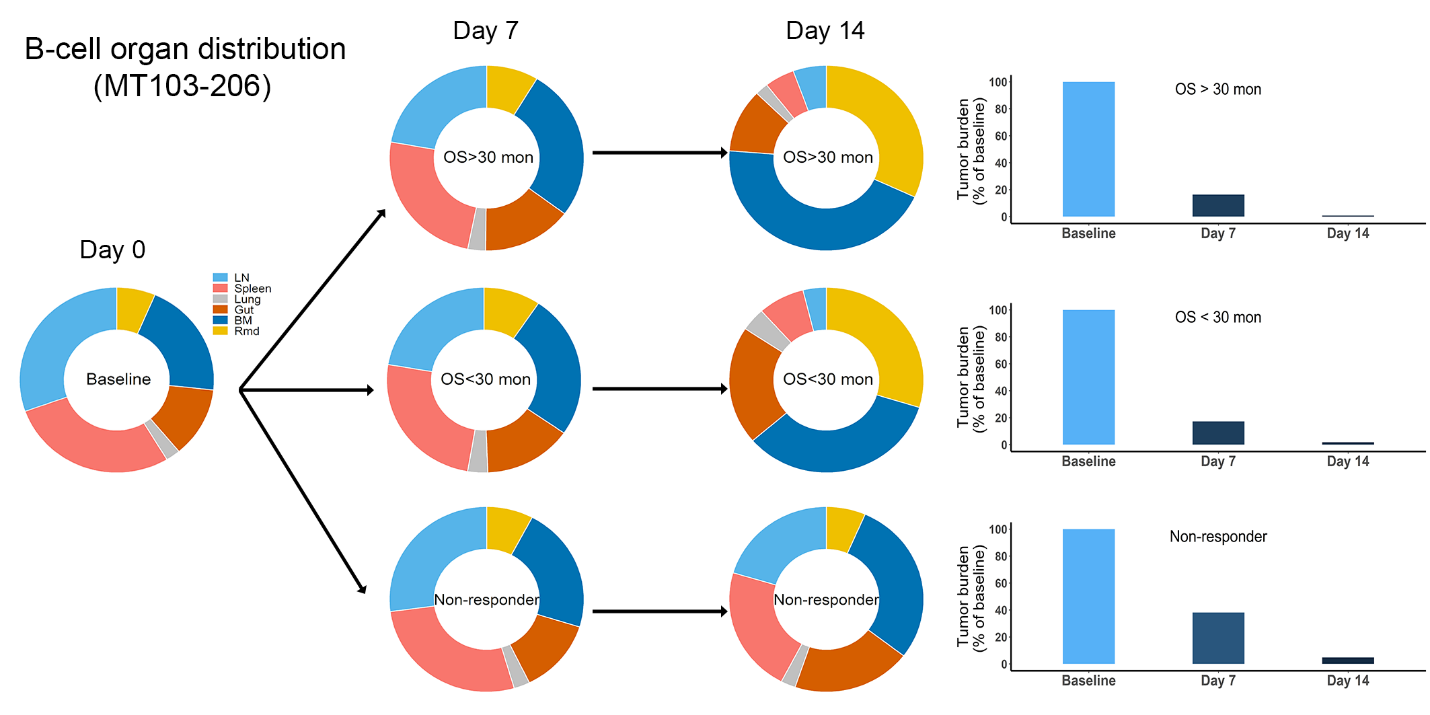

Figure S16 Simulated baseline (Day 0), and post-treatment (Day 7 and 14) B-cell organ distribution in patients of MT103-206. The in-vivo model was applied to perform the simulations. Bar plot showed the simulated tumor burden at baseline and post-treatment. OS, overall survival.

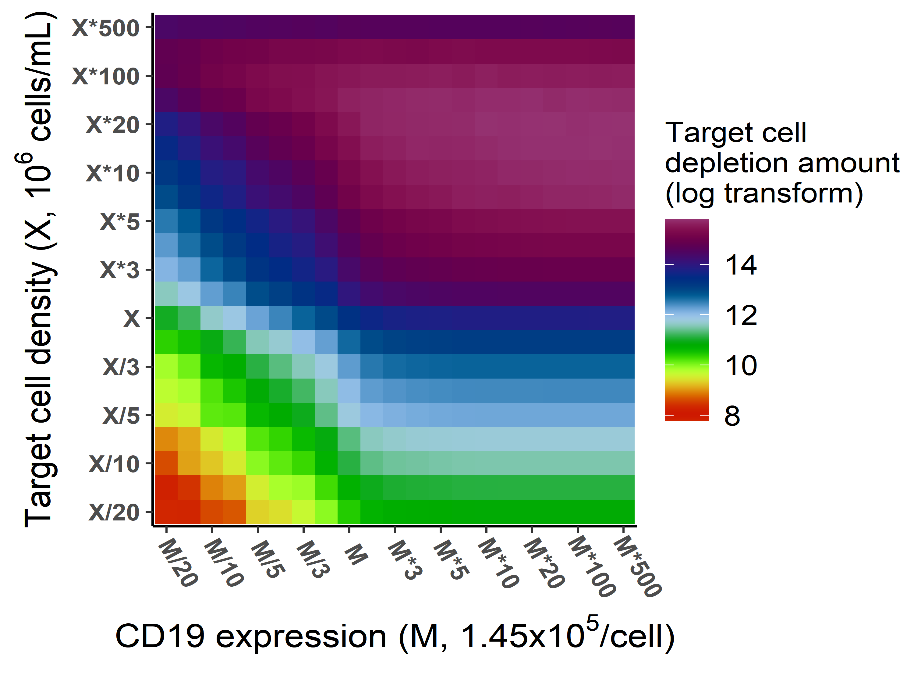

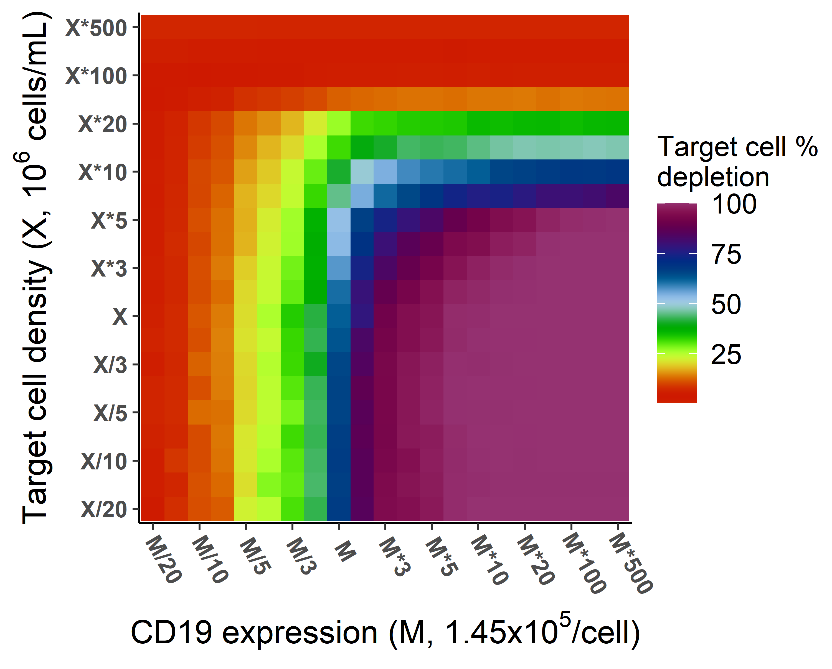
**a	b**

A (M,X)

B (M,10 X)

C (10M, X)

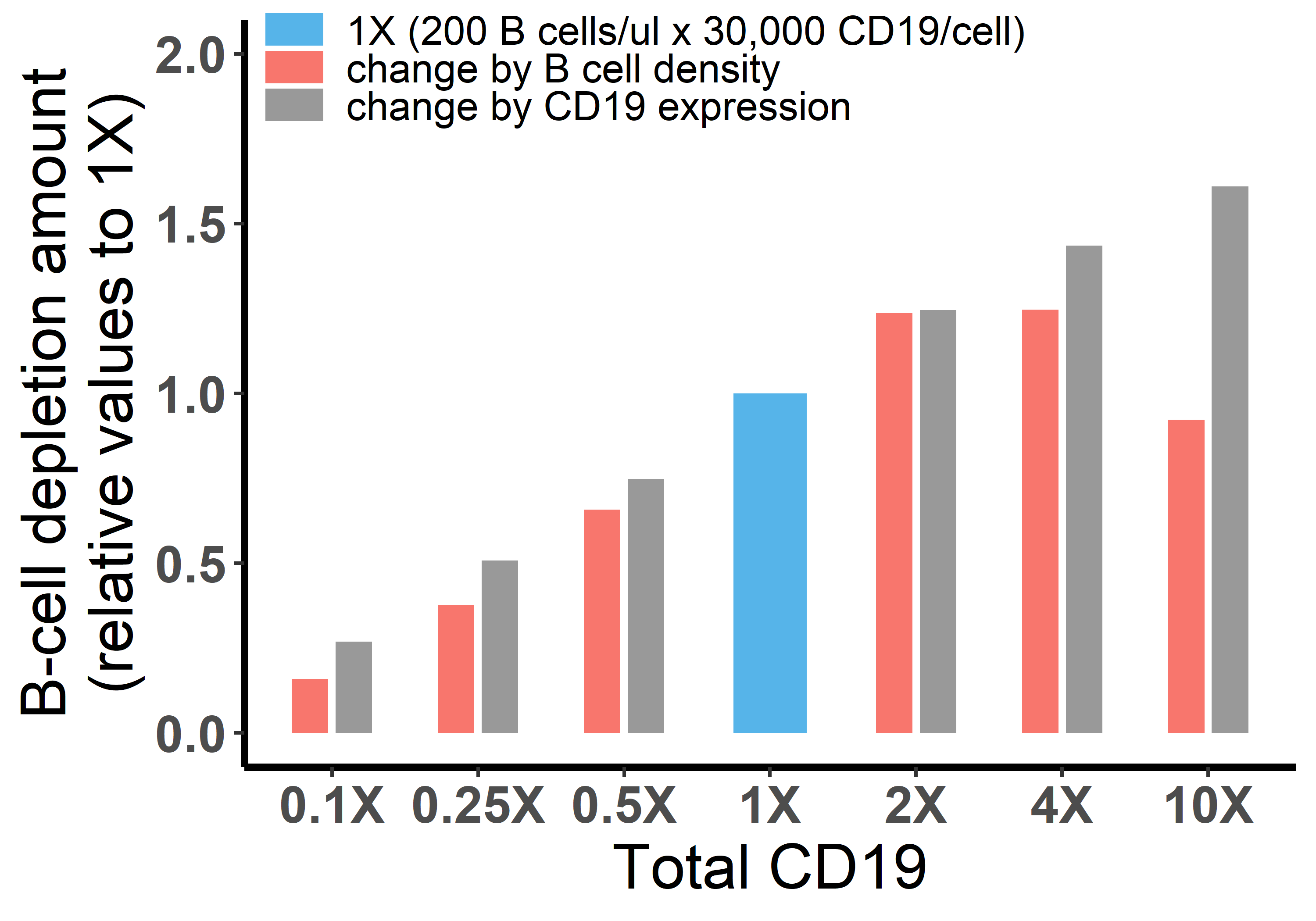
**c**

Figure S17 Different effects of CD19 on cellular and molecular processes. **a-b,** Different effects of target cell density and CD19 expression on target cell depletion amount (**a**) and fraction (**b**). The in-vitro model was used in the simulation. Initial setup: CD3(L), effector cell (1X), target cell (X/20 to 500X), CD19 expression (M/20 to 500M), 0.65 ng/mL, 72h.

Total CD19 in the system was jointly influenced by CD19 expression per cell and target cell density. CD19 expression influenced cell lysis to a similar extent as target cell density when both factors were low (e.g. CD19 expression < M, target cell density < X in Fig. S17a). However, further increase of CD19 expression on cell membrane did not further improve cell lysis (e.g. from point A to point C), indicating maximum ternary complexes at each interface has been reached. In contrast, the increase of target cell density continuously to promote cell lysis, e.g. from point A to B, due to enhanced probability of cell-cell encounter. When the target cell density reaches extremely high (> 50 X), cell lysis started to decrease, resulting from fewer cell-cell adhesion events due to insufficient BiTE concentration. As shown, although total CD19 in the system at point B and C are identical (10·M·X), different cell lysis level is yielded, supporting different effects of CD19 on cellular and molecular processes.

**c,** Effects of B cell density and CD19 expression on B cell depletion in vivo. In each group, the change of total CD19 density from the reference (1X, 200 B cells/μL × 30,000 CD19/cell) was achieved through changing B cell density (red bar) or CD19 expression (grey bar). Their effects on B cell depletion amount were simulated by the in-vivo model. Initial setup for reference (1X): T cell (200/μL), B cell (200/μL), CD3 (50, 000/cell), CD19 (30, 000/cell), 0.73 ng/mL, 72h.

Different effects of CD19 on cellular and molecular processes have been confirmed by in-vivo model. Similarly, the increase of CD19 expression within low level range (3×10^3^ to 3×10^5^ CD19/cell from 0.1X to 10X) constantly improved cell depletion, as ternary complexes formation at each interface increased with CD 19 expression. By contrast, bidirectional effect was shown by increasing B cell density, owing to enhanced probability of cell-cell encounter and then insufficient BiTE concentration. Herein, B cell density at 10X (2,000/μL) in blood indicates extremely high organ B cell density in the model, e.g. spleen (~ 9×10^8^/mL) and lymph nodes (~ 3×10^8^/mL).

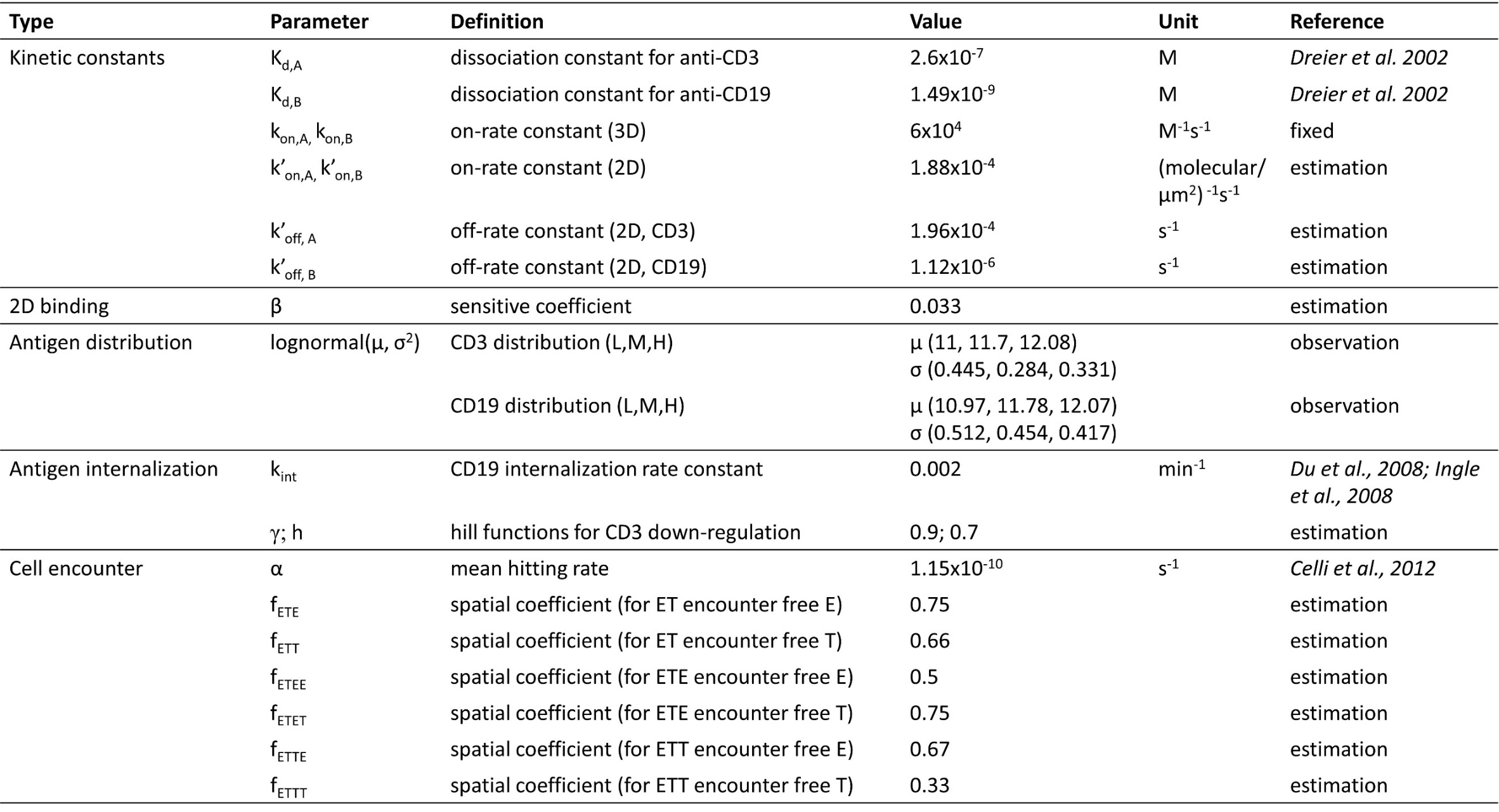
Table S1 Important model parameters (for base model)

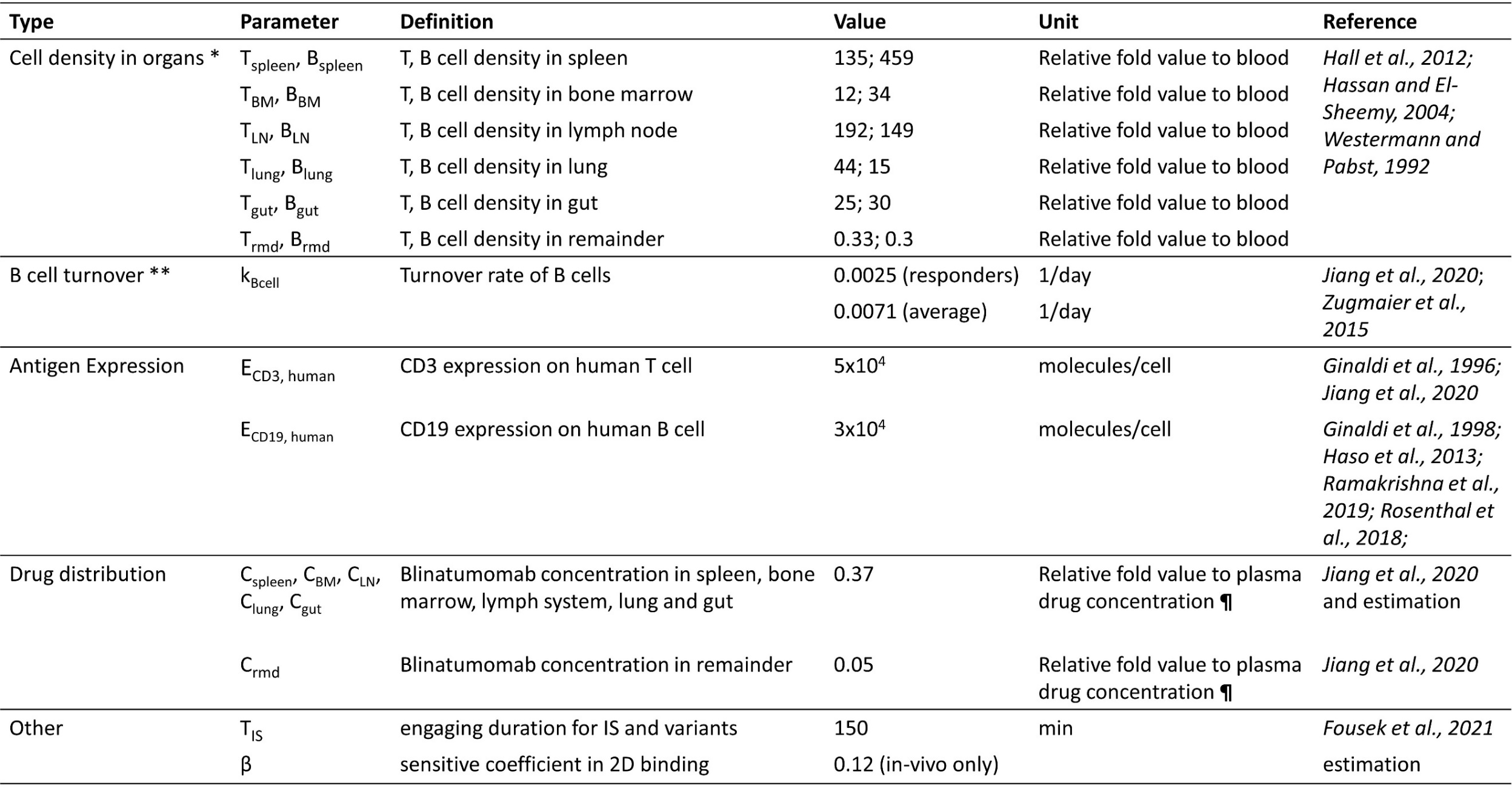
Table S2 Patient specific parameters for the in-vivo model

* T cell density in organ during treatment was allowed to change over time in the simulation. The values were derived from the change of blood T cell count based on reported data (***Bargou et al. 2008; Zhu et al., 2016; Zhu et al., 2018;*** [***Zugmaier***](https://pubmed.ncbi.nlm.nih.gov/?term=Zugmaier+G&cauthor_id=26480933) ***et al., 2015***). The blood T cell decline due to rapid redistribution after administration was excluded.

** In the simulation, the value of 0.0025/day was applied to “Responder” group in trial MT103-211 and “OS>30 month” and “OS<30 month” groups in trial MT103-206; the value of 0.0071/day was applied to MT103-202, and “Non-responder” group in trial MT103-206. B cell turnover rates for trial MT103-104 and “Non-responder” group in trial MT03-211 were estimated as 0.0067/day and 0.1/day respectively.

**¶** plasma drug concentration is trial-specific, see ***Supplementary Table S3***.

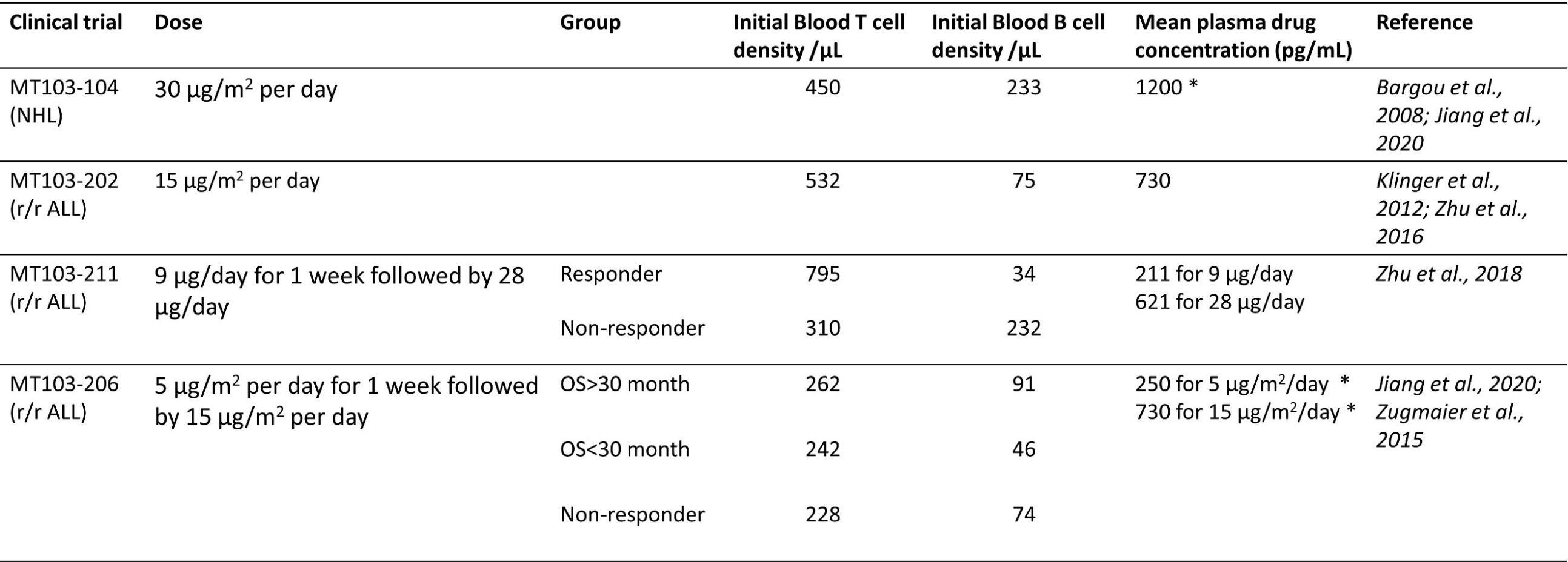
 Table S3 Clinical trial information (baseline cell density in blood and mean plasma drug concentration at steady state)

NHL, non-Hodgkin's Lymphoma; OS, overall survival; r/r ALL, relapsed/refractory acute lymphoblastic leukemia.

* For MT103-104 and MT103-206, mean plasma drug concentration was derived from a linear relationship between dose and blood drug concentration (***Jiang et al., 2020***).
